# Fine-scale Population Structure of North American *Arabidopsis thaliana* Reveals Multiple Sources of Introduction from Across Eurasia

**DOI:** 10.1101/2021.01.22.427575

**Authors:** Gautam Shirsekar, Jane Devos, Sergio M. Latorre, Andreas Blaha, Maique Queiroz Dias, Alba González Hernando, Derek S. Lundberg, Hernán A. Burbano, Charles B. Fenster, Detlef Weigel

**Affiliations:** Max Planck Institute for Developmental Biology, 72076 Tübingen, Germany; Centre for Life’s Origin and Evolution, University College London, London WC1E 6BT, UK; South Dakota State University, Brookings, SD 57007, USA

**Author notes:** Centre for Life’s Origin and Evolution, University College London, London WC1E 6BT, UK (S.M.L.); Federal University of Viçosa, 36570-900 Viçosa - MG, Brazil (M.Q.D.).

**Keywords:** *Arabidopsis thaliana*, population genetics, admixture, non-native species, migration

## Abstract

Large-scale movement of organisms across their habitable range, or migration, is an important evolutionary process that can contribute to observed patterns of genetic diversity and our understanding of the adaptive spread of alleles. While human migrations have been studied in great detail with modern and ancient genomes, recent anthropogenic influence on reducing the biogeographical constraints on the migration of non-native species has presented opportunities in several study systems to ask the questions about how repeated introductions shape genetic diversity in the introduced range. We present here the most comprehensive view of population structure of North American *Arabidopsis thaliana* by studying a set of 500 (whole-genome sequenced) and over 2800 (RAD-seq genotyped) individuals in the context of global diversity represented by Afro-Eurasian genomes. We use haplotype-sharing, phylogenetic modeling and rare-allele sharing based methods to identify putative sources of introductions of extant N. American *A. thaliana* from the native range of Afro-Eurasia. We find evidence of admixture among the introduced lineages that has resulted in the increased haplotype diversity and reduced mutational load. Further, we also present signals of selection in the immune-system related genes that impart qualitative disease resistance to pathogens of bacterial and oomycete origins. Thus, multiple introductions to a non-native range can quickly increase adaptive potential of a colonizing species by increasing haplotypic diversity through admixture. The results presented here lay the foundation for further investigations into the functional significance of admixture.

## Introduction

When an organism is introduced outside its native range where it has established its eco-evolutionary history, how well the organism adapts to the new environment depends on diverse factors, including history of introduction, founder effects, and natural selection. Such factors are crucial to our understanding of the genetic changes associated with adaptation in introduced lineages (Colautti and Lau 2015; Estoup et al. 2016). North America’s post-Columbian human colonization has facilitated mostly unidirectional cross-continental species movement that includes several plant species (La Sorte, Mckinney, and Pyšek 2007; Winter et al. 2010), and presents a unique natural experiment to study the role of genetic history in explaining extant plant diversity and its impact on plant adaptation.

Our understanding of human colonization history of North America, commonly referred to as “peopling” of America, has been greatly advanced through genetic and/or archaeological evidence from the pre-Columbian era (Reich et al. 2012; Skoglund et al. 2015; Potter et al. 2018; Flegontov et al. 2019; Becerra-Valdivia and Higham 2020). Recent studies on post-Columbian impact on current human population structure have taught us the complex effects of global migrations on human genetic diversity in N. America (Bryc et al. 2015; Han et al. 2017; Dai et al. 2020). Inadvertently, humans have also introduced many commensal species to N. America, and these can potentially provide informative complements and contrasts to demographic processes during colonization observed in humans (La Sorte, Mckinney, and Pyšek 2007). Specifically, important questions are how much of the native diversity was introduced to N. America, how much new diversity has been generated in situ through mixing of lineages that originated from distant parts in the native range, and how much the observed diversity has been shaped by selection.

*Arabidopsis thaliana* is a commensal of humans that is native to Africa and Eurasia (Fulgione and Hancock 2018). Being a model species for plant research, it has an extensive list of resources that include range-wide whole genome sequences (1001 Genomes Consortium 2016; Durvasula et al. 2017; Zou et al. 2017; Hsu, Lo, and Lee 2019) and genome annotation built on decades of rigorous molecular biology research that allows one to explore genetic history of the organism in detail. In the post-Columbian era, *A. thaliana* has migrated to N. America, almost certainly enabled by humans, and has established itself on a wide geographic range across the continent. Coarse-scale population structure analysis of N. American individuals with 149 single nucleotide polymorphism (SNP) markers has revealed the presence of a dominant lineage “Haplogroup1” (Hpg1) (Platt et al. 2010). Patterns of mutation accumulation in the genomes of Hpg1 individuals have supported an arrival in N. America about 400 years ago, early after Europeans started to arrive en masse on the continent. A parsimonious explanation of the ubiquitous nature of this lineage could be that it was the earliest to be introduced to N. America (Exposito-Alonso et al. 2018). So far, little consideration has been given to the supposedly later arrival of other lineages, their origins in the native range, their fate as migrations continued during the past centuries and how their genomes have been shaped by processes such as admixture and adaptation.

We present the fine-scale population structure of the North American *A. thaliana* population as seen through the lens of range-wide genetic diversity of the species. Using genomes of *A. thaliana* individuals collected from the Midwest, the Eastern Seaboard and the North-East of the current United States of America, we infer possible sources of ancestry based on haplotype-sharing, phylogenetic tree-based modeling and rare allele sharing with the worldwide dataset. We also describe how admixture in this predominantly selfing species is generating new haplotype diversity and how admixture affects the fate of deleterious mutations and allows selection on immunity related loci. Our findings highlight how tools developed for the genetic study of human migrations and ancestry can be productively applied to other species in order to learn about their recent history. Most importantly, the work presented here shows that increased global connectivity through the past two centuries has made species invasions from across the species range common.

## Results

### An overview of population structure and genetic drift from RAD-seq

We collected *A. thaliana* samples across an area of about 1,200 by 900 km in the Eastern United States of America during the spring seasons (mid-March to early June) of 2014, 2015 and 2016 (Fig. 1; Table S1a,b). We genotyped these samples using a RAD-seq implementation of reduced representation sequencing (Miller et al. 2007). After filtering for sequencing output and quality, we retained 2,861 individuals, which shared 4,907 polymorphic SNPs. In order to compare the population structure and genetic diversity in our N. American to the global Afro-Eurasian collection (AEA) we used data from the 1001 Genomes project (1001 Genomes Consortium 2016) in addition to whole genome sequences from 13 Irish (this work), 10 African (Durvasula et al. 2017) and 5 Yangtze River basin accessions (Zou et al. 2017). From these AEA individuals, information on the 4,907 polymorphic SNPs found in our N. American individuals (average depth ~36X) were extracted and merged with the N. American dataset for further analysis.

**Figure 1.**
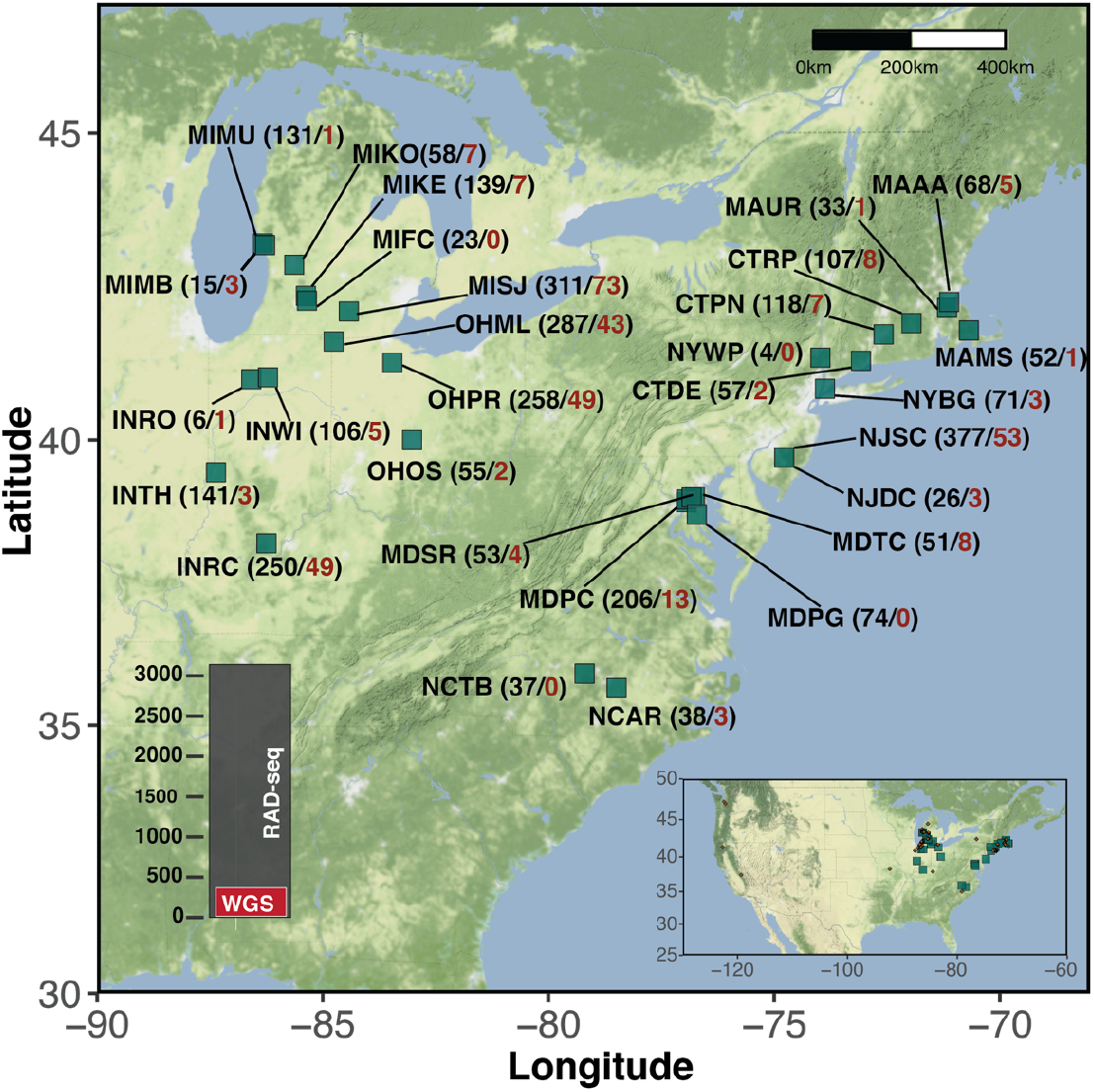
Locations and number of sampled individuals. Abbreviations of the locations sampled are shown along with the number of RAD-sequenced samples (in black) and the number of whole-genome sequenced (WGS) samples (in red). Left inset: bar plot of total number of samples sequenced. Right inset: sampling area in the context of N. America.

Although pairwise similarity using “identity-by-state” (IBS) and “identity-by-descent” (IBD) across the genome is greater in N. American than in AEA individuals, genetic drift relative to AEA individuals could nevertheless be observed in N. American individuals with principal component analysis (PCA) (Fig. S1A). North American individuals in our collection were genetically much more diverse than the N. American individuals previously sequenced as part of the 1001 Genomes Project (Fig. S1B).

### Diversity of N. American haplogroups

We first used RAD-seq to rapidly genotype thousands of individuals, but because of its inherent biases (low density of markers, strand-bias, underestimation of genetic diversity), these data are not well suited for fine-scale, quantitative population genomic analyses (Cariou, Duret, and Charlat 2016; B. Arnold et al. 2013; Lowry et al. 2017). We therefore selected a subset of distantly related individuals for whole-genome sequencing, at an average of ~ 8x coverage. A PCA of 500 N. American individuals, including a subset of previously analyzed herbarium individuals (Exposito-Alonso et al. 2018) (Fig. S1C), resulted in an arrangement in which most individuals were found along distinct clines. We decided to explore this population structure in detail using different complementary population genetic methods.

Finer-scale population structure can be revealed by explicitly modeling the effects of linkage disequilibrium (LD) and clustering individuals based on their shared ancestry that emerges after accounting for LD (Montinaro et al. 2015; Busby et al. 2015; Leslie et al. 2015). Therefore, we hierarchically partitioned the N. American individuals into 57 clusters (from here on called *groups*) using a co-ancestry matrix derived using CHROMOPAINTER v2 and MCMC-based clustering in fineSTRUCTURE (Lawson et al. 2012) (Fig. 2A, B). Haplogroup1 (Hpg1) is the most frequently observed *group* across the sampled populations (Fig. S3), consistent with previous observations (Platt et al. 2010; Exposito-Alonso et al. 2018). OHML (Ohio) and NJSC (New Jersey) had the highest within-population haplotype diversity, with 11 and 12 *groups*. Several *groups*, such as OhioNewJersey2, IndianaNewJersey1 and NewJerMich1, were found in populations from geographically distinct regions (Fig. S3).

**Fig 2.**
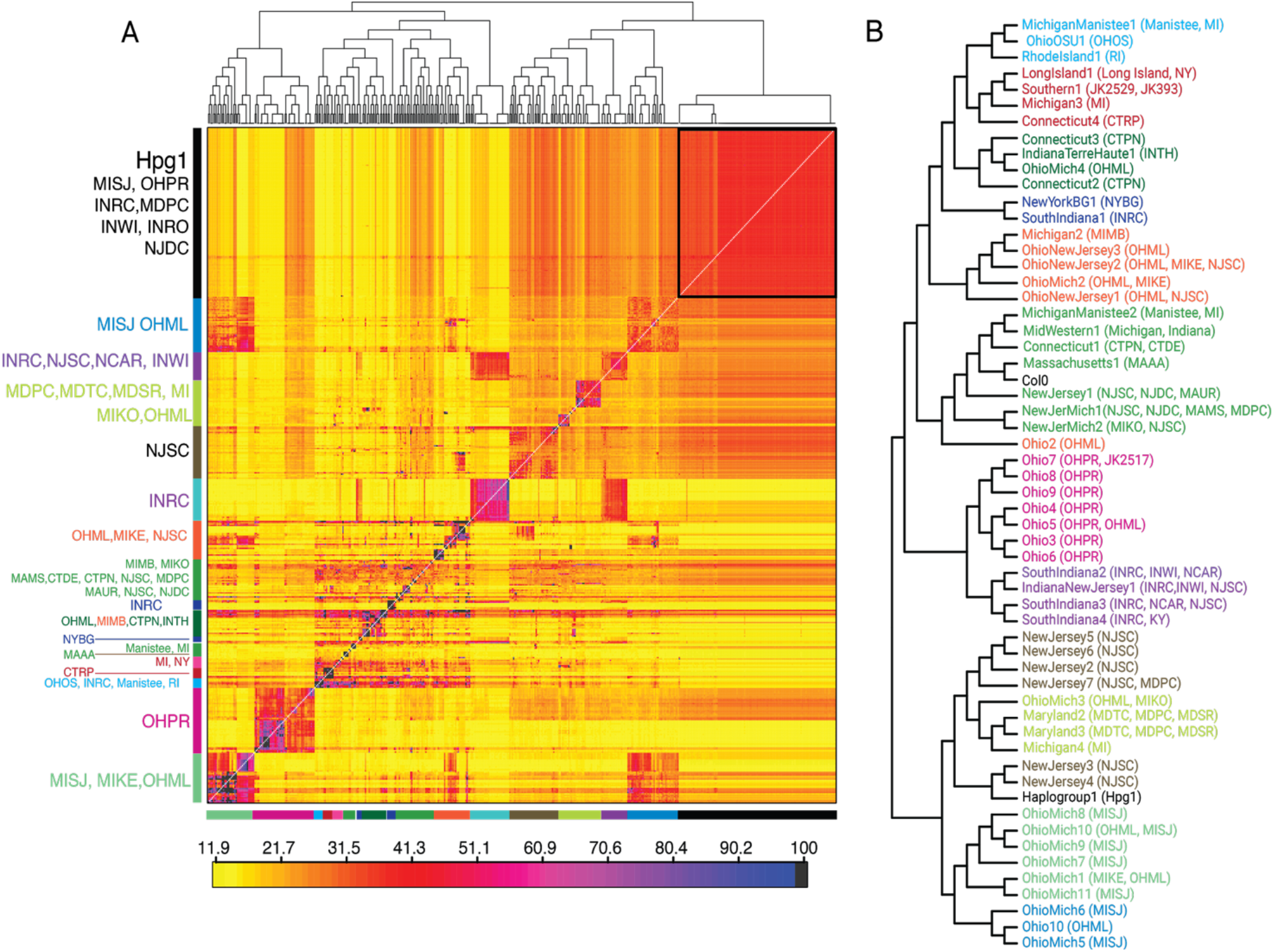
Identification of different haplogroups in N. American individuals. A. Co-ancestry matrix generated by chromosome painting of each N. American individual against all others (using CHROMOPAINTER) and ordered by FINESTRUCTURE analysis, B. Collapsed FINESTRUCTURE tree generated by merging individuals into groups (populations from which the individuals were collected are shown in parentheses, herbarium individuals are denoted by JKxxx).

We further analyzed the genetic relationships among these *groups* using several complementary approaches. Treemix (Pickrell and Pritchard 2012), without considering migration edges, reconstructed relationships among the *groups* (Fig. S4A) similar to the topology inferred by fineSTRUCTURE clustering (Fig. 2B). Residuals from the fitted model indicated higher genetic relatedness within populations (positive high values), but also pointed towards probable gene flow from haplogroup1 to some of the other *groups*. That Hpg1 has contributed to these new *groups* is consistent with Hpg1 being the most frequent *group* and having the widest geographical distribution.

Stochastic changes in allele frequency, as a result of the neutral process of drift, hold information about shared ancestry. We therefore estimated values for the outgroup *f*_*3*_ statistic (Patterson et al. 2012) to understand the shared drift among *groups* relative to an outgroup. Indeed, some of the N. American groups (OhioMich1, SouthIndiana4 and Ohio7) along with Hpg1 shared excess drift with other *groups* (Fig. S5), pointing to these *groups* as a putative source of gene flow. Next, we calculated values for the *f*_*3*_ statistic in all trios (*groupA*, *groupB*: *groupTest*) of N. American *groups* to detect whether *groupTest* was admixed between *groupA* and *groupB*. There were several *groupTest* examples with negative f3 scores and *Z-*scores below −3 in several trios (Fig. S6A). In several cases, Hpg1 emerged as a putative source (as either *groupA* or *groupB*) (Table S2). To investigate this in more detail, we calculated the shared drift of Hpg1 relative to the other N. American *groups*. We found more *groups* with high levels of shared drift with Hpg1 than *groups* with limited shared drift; Massachusetts1 and MichiganManistee1 were the *groups* with least shared drift (Fig. S6B). We calculated the ABBA-BABA statistic (D-statistic) in the form of (*Massachusetts1, Test: Haplogroup1, MichiganManistee1*) to learn the extent of gene flow between Hpg1 and other N. American *groups* (FigS6C). Many *groups* showed significantly more ABBA sites (Z-score <−3) than BABA sites, confirming the contribution of Hpg1 ancestry to the genetic makeup of these *groups*.

### Contribution of distinct sources of ancestry to N. American diversity

It is clear from the above that there must have been more than one introduction of *A. thaliana* to N. America. What is not clear is whether the observed haplogroups already existed in Eurasia, or whether they only formed by intercrossing in N. America. We therefore wanted to learn whether N. American extant haplogroups include ancestry from different geographic regions in Eurasia. We first excluded lineages that showed evidence of recent admixture (*groups* with significantly negative *f*3*-* scores), and we then applied statistical procedures based on shared haplotype chunks (fineSTRUCTURE), shared drift (outgroup *f*_*3*_, D-statistic and qpWave) and enrichment of rare alleles with respect to the AEA haplotype diversity to identify sources of ancestry in Eurasia based on whole genome sequences from AEA individuals (n=1039) (1001 Genomes Consortium 2016).

We traversed the genomes of N. American individuals to assign local ancestry along each chromosome. To this end, we performed haplotype based inference in three steps: (i) Paint each AEA individual against the others (excluding itself) with CHROMOPAINTER v2, (ii) Based on haplotype sharing, cluster individuals into *sub-clusters* using fineSTRUCTURE. These “sub-clusters” were then grouped into *clusters*, and *clusters* were further grouped into *regions* (Fig. 3A,B; details of these hierarchical partitions for each AEA individual are given in Table S3). (iii) Estimate an ancestry profile for individual N. American groups (recipients) as a mosaic of AEA donors from different *regions*.

**Fig. 3.**
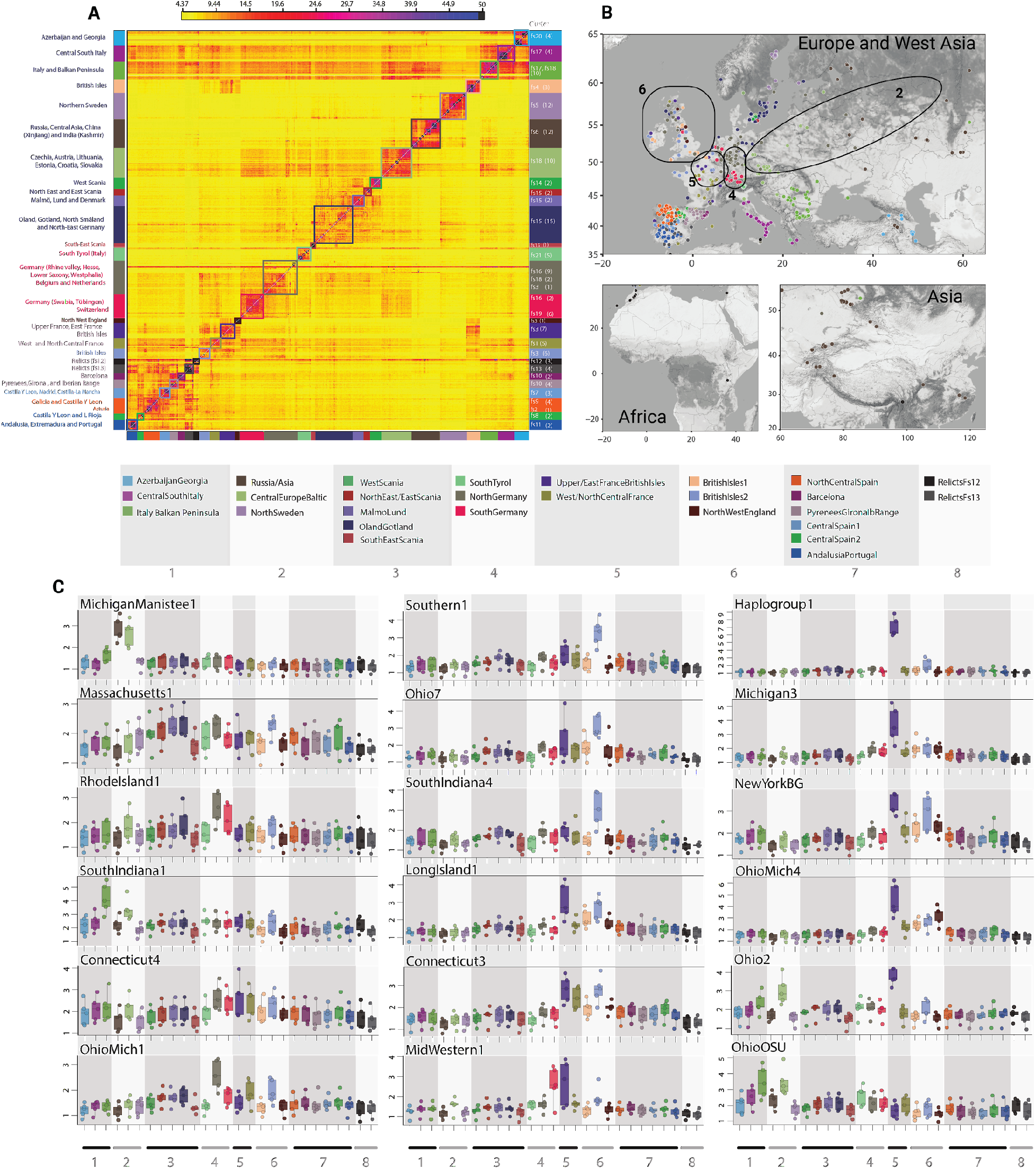
fineSTRUCTURE clustering of non-American *A. thaliana* individuals from the native range and chromosome painting of N. American groups with the *regions* as donors. **A.** Co-ancestry matrix of *A. thaliana* derived by chromosome painting of individuals from the 1001 Genomes project (excluding N. American individuals) and additional genomes from China, Ireland and Africa (using CHROMOPAINTER). Subsequent fineSTRUCTURE clustering resulted in 21 *regions* and 157 sub-clusters. **B.** Geographic locations of individuals from (A) colored by their membership in one of the 21 *regions*. Numbered *regions* (2, 4, 5, 6) marked with black boundaries are main sources of introduction. **C.** Chromosome painting of N. American groups using 157 *sub-clusters* (normalized by number of donor individuals per *sub-cluster* and averaged to per chromosome per *region*). Numbering of *regions* 1-8 according to bottom of (B).

Box plots in Fig. 3C show these inferred ancestry profiles for the N. American *groups*. It can be seen that although the majority of groups are enriched for Upper/EastFranceBritishIsles ancestry, other British Isles *regions* (BritishIsles1, BritishIsles2 and NorthWestEngland) also feature significantly across several groups. Apart from these, some N. American *groups* such as MichiganManistee1, Ohio OSU, Ohio2, SouthIndiana1 had substantially higher contributions from East European *regions* such as RussiaAsia, CentralEurope/Baltic and Italy/BalkanPeninsula. NorthGermany and SouthGermany *regions* have contributed to the ancestry of OhioMich1, RhodeIsland1 and Mid-Western1 *groups* (Fig3C).

We explored these haplotype sharing patterns further by measuring shared drift between a test N. American *group* and 158 *sub-clusters* of AEA individuals using outgroup *f*_*3*_ statistic of the form *test, sub-cluster*; relictsFs12-3 (We chose relictsFs12-3 as an outgroup as it is a highly diverged *sub-cluster* comprising relict population individuals). At a coarser scale, the results agree with the haplotype-based inferences. Shared allelic drift measured with outgroup *f*_*3*_ statistic showed that the current N. American *groups* are related to the AEA *sub-clusters* that belonged to either western, central or eastern Europe (Fig. S7). We also observed these patterns of relatedness qualitatively in a PCA plot where we projected N. American individuals into PC space occupied by AEA individuals (Fig. 4A). Even finer details became apparent with uniform manifold approximation and projection (UMAP) embeddings (McInnes, Healy, and Melville 2018) (Fig. 4B) derived from the first 50 PC components of all the individuals (without projection).

**Fig. 4.**
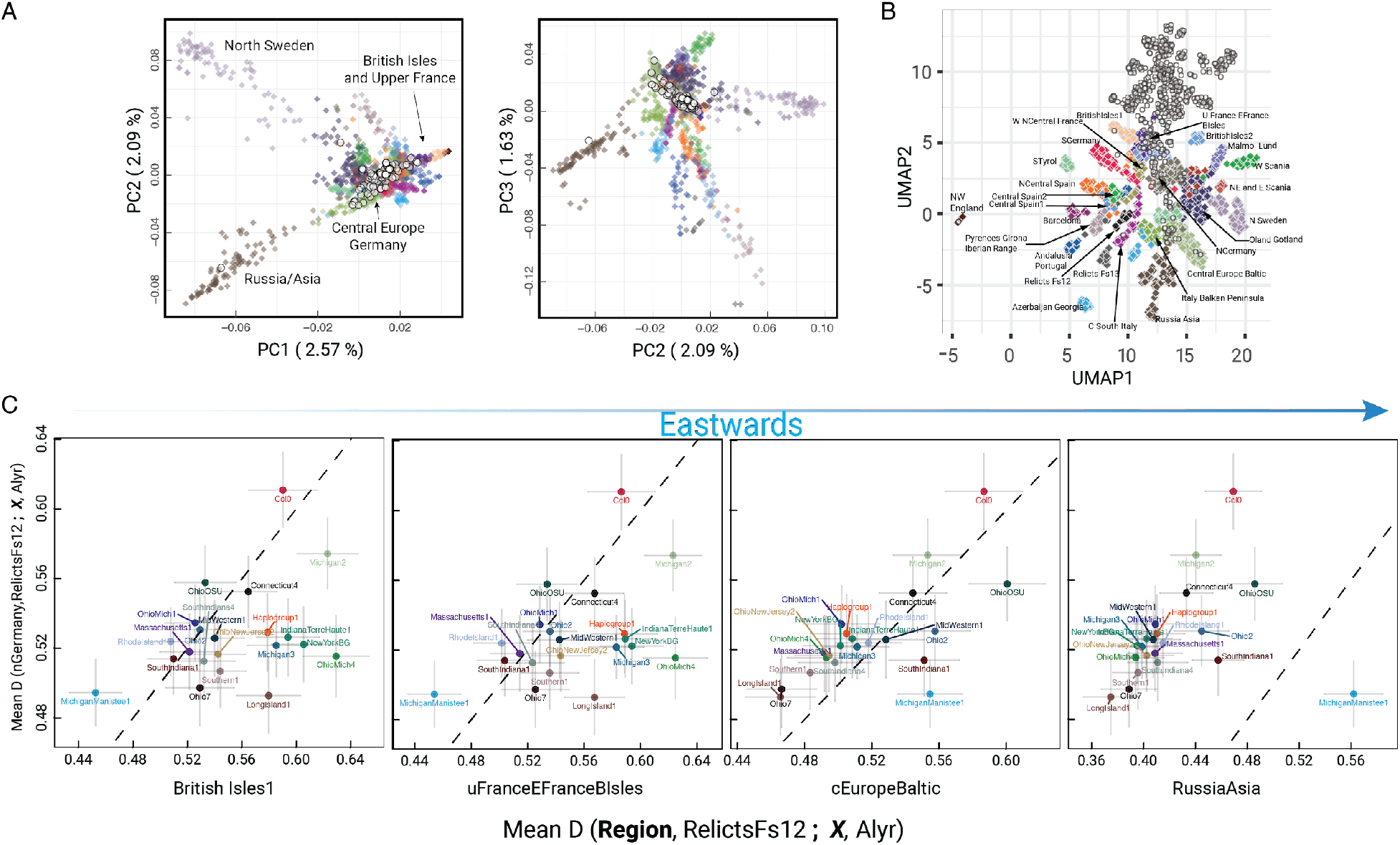
Multiple sources of origin of N. American haplogroups. **A.** Projection of N. American individuals (white circles) in PC space derived from individuals of 27 Afro-Eurasian regional clusters. **B.** Uniform manifold approximation and projection (UMAP) embeddings of the first 50 PC components derived from PCA (without projection for N. American individuals) of all individuals. (White circles: N. American individuals). **C.** Biplot of mean D-statistics of N. American haplogroups (X) with *sub-clusters* comprising NorthGermany region (*NorthGermany*, *RelictsFs12*; *X*, Alyr), against mean *D*-statistics of *sub-clusters* comprising different *regions* in an eastward direction (*testRegion*, RelictsFs12; ***X***, alyr). Vertical and horizontal bars represent the spread of D-statistics from member *sub-clusters* of each *region*.

The coarse patterns of shared ancestry emerging from outgroup *f*_*3*_ statistic, PCA projection, and UMAP embeddings were tested in a more systematic way by modeling phylogenies of specific topographies. We first calculated *D*-statistic (Patterson et al. 2012) for all the N. American non-admixed *groups* (X) in the form of (NorthGermany, RelictsFs12; X, alyr). NorthGermany was chosen because of its central location among the possible sources of N. American *A. thaliana*. We then calculated *D-* statistics for the N. American *groups* by replacing NorthGermany with 4 *regions*: BritishIsles1, Upper/EastFranceBritishIsles, CentralEurope/Baltic and RussiaAsia. We then plotted *D*-statistics with replacement *regions* to *D*-statistics obtained using NorthGermany separately. These biplots (Fig. 4C) clearly differentiate western and eastern European sources of ancestry in N. American *A. thaliana*. OhioOSU, Ohio2, SouthIndiana1 and MichiganManistee1 clearly showed the relative eastern European ancestry component. The analysis also revealed that Col-0, the reference genome accession for *A. thaliana* research, shares significant ancestry with individuals from NorthGermany, confirming the origin of Col-0 in or near Germany (Rédei 1992).

We extended this analysis using qpWave (Reich et al. 2012) to test whether any two N. American *groups* would be symmetrically related to a set of outgroups (AEA *regions*). Specifically, we tested whether a set of *f*_*4*_-statistics comprising two N. American *groups* across a set of layer1 outgroups (AzerbaijanGeorgia, Barcelona, NorthSweden, NorthWestEngland, Relicts Fs13, SouthTyrol, WestScania and West/NorthCentralFrance) makes a matrix of rank 0 (same wave of ancestry) (Table S4). We then tested whether addition of an extra outgroup region (consisting of putative sources of ancestry) to the layer 1 outgroup set affected the symmetry of shared ancestry. If the two test N. American *groups* are differentially related to the extra outgroup region, then it would increase the rank of the original matrix of *f*_*4*_-statistics (rejection of rank 0), indicating distinct streams of the ancestry among the test *groups*. We added an extra outgroup from additional regions of BritishIsles2, ItalyBalkanPeninsula, NorthGermany, RussiaAsia and Upper/EastFranceBritishIsles one-by-one. Adding these putative source regions affected the symmetrical relationships observed with our original outgroup set. Except in the case of SouthIndiana4 and Ohio7, all the N. American *group* combinations showed asymmetric relationships (rejection of rank 0) with these extra outgroups (Table S4).

These results validated the findings from qualitative observations made with outgroup *f*_*3*_ statistic, PCA projection, and UMAP embeddings. It further confirmed results obtained from *D-*statistics analysis, that the N. American *A. thaliana groups* have ancestral components from western Europe (mainly British Isles), central Europe and eastern Europe.

More subtle patterns of ancestry can be inferred by finding rare variants from AEA that have risen to higher frequency in N. American individuals. Because we had moderate- to high-coverage whole genomes of the AEA and N. American individuals, we could use such rare variants to independently ascertain the results obtained from the haplotype-based ancestry inference and shared ancestry based inference, mostly on moderate to high frequency alleles. We identified variants from AEA individuals with frequency of 1% or lower and tracked their enrichment in the N. American *groups*. We found that different N. American *groups* have accumulated rare alleles from different AEA *sub-clusters* (Fig. S8). Whereas several N. American *groups* have inherited rare alleles from British Isles *sub-clusters*, *groups* RhodeIsland1, MichiganManistee1, OhioOSU, SouthIndiana1 and OhioMich1 have accumulated rare alleles from central/eastern Europe sub-clusters, while Hpg1 has accumulated a significant number of rare alleles from *sub-clusters* from the Upper/EastFrance/BritishIsles *region*. Taken together, this analysis confirmed that N. America was colonized by *A. thaliana* in multiple waves with distinct sources of ancestry.

### Environmental conditions at source and success of colonizing lineages

As we had inferred the shared ancestry of the colonizing lineages with different complementary methods, we hypothesized that besides human-assisted migration, environmental similarity between putative source *sub-clusters* and colonizing lineages contributed to successful colonization of the lineages. To test this hypothesis, we fit a regression model to predict shared ancestry with AEA *sub-clusters* (measured by outgroup *f*_*3*_ statistics of the form test *N. American group, AEA sub-cluster: RelictsFs12_3 (outgroup)*), using linear combinations of four environmental variables: average temperature (tavg), precipitation (prec), solar radiation (srad) and water vapor pressure (vapr) in a Bayesian multilevel modeling (bMLM) framework (Gelman 2006). We used the bMLM strategy in order to understand each N. American group’s environmental association with its putative source AEA *sub-clusters* without ignoring the environmental association to the entire cohort of N. American *groups*.

Population-scale coefficients for the environmental variables precipitation (mm) and water vapor pressure (kPa) revealed that environmental dissimilarity calculated by Euclidean distance between each N. American *group* and AEA *sub-cluster* is negatively correlated with the outgroup *f*_*3*_ statistics (Table 1). Although average temperature dissimilarity is slightly negatively correlated with outgroup *f*_*3*_ statistics, the compatibility interval with the model is large, with slightly positive correlation in posterior distribution. Upon closer examination of the coefficients estimated for individual N. American *groups*, it can be seen that precipitation and water vapor pressure dissimilarity is negatively correlated with the outgroup *f*_*3*_ statistic for all *groups* but MichiganManistee1 (Fig. S9). Overall the general trend of negative correlation of the linear combination of the dissimilarity of the variables (average temperature, precipitation, solar radiation, and vapor pressure) to the outgroup *f*_*3*_ statistic can be captured with the individual estimates sampled from the posterior distribution (Fig. S10).

**Table 1.** Posterior summary of the regression coefficients for environmental variables. Bayesian multilevel model based pooled estimates of regression coefficients for environmental variables T_avg_ (°C), Precipitation (mm), solar radiation (kJ m^−2^ day^−1^) and water vapor pressure (kPa). Outgroup f_3_ statistic of each N. American group to every AEA sub-cluster (outgroup: relicts Fs12_3) was used as a Student’s t-distributed dependent variable.

| Parameter | mean | sd | hdi_3% | hdi_97% | $\hat{R}$ |
| --- | --- | --- | --- | --- | --- |
| $\bar{a}$ | 0.1390 | 0.0920 | -0.0280 | 0.3190 | 1.0000 |
| $\bar{\beta}_{T_{avg}}$ | -0.0640 | 0.0960 | -0.2420 | 0.1170 | 1.0000 |
| $\bar{\beta}_{Prec}$ | -0.1800 | 0.0910 | -0.3530 | -0.0080 | 1.0000 |
| $\bar{\beta}_{Srad}$ | -0.0010 | 0.0910 | -0.1800 | 0.1600 | 1.0000 |
| $\bar{\beta}_{V_{apr}}$ | -0.2320 | 0.0960 | -0.4080 | -0.0490 | 1.0000 |
| $\sigma$ | 0.3950 | 0.0110 | 0.3730 | 0.4160 | 1.0000 |

The negative correlation between environmental dissimilarity and shared ancestry led us to hypothesize that in reduced dimensional space of environmental variables (average temperature, precipitation, and vapor pressure), N. American *groups* should occupy space near their source AEA *sub-clusters*. To test this, we performed UMAP on the standardized values for environmental variables for N. American *groups* and AEA *clusters* together. We observed that the N. American groups and their putative source *clusters*, as inferred by population genomic approaches (specifically *sub-clusters* from Upper/EastFranceBritishIsles, NorthGermany, SouthGermany, BritishIsles1, BritishIsles2 and CentralEurope/Baltic *regions*) occupied similar space in the UMAP embeddings (Fig. 5), thus confirming that overall environmental similarity between source populations and N. America might be an important contributor to the success of colonization.

**Fig. 5.**
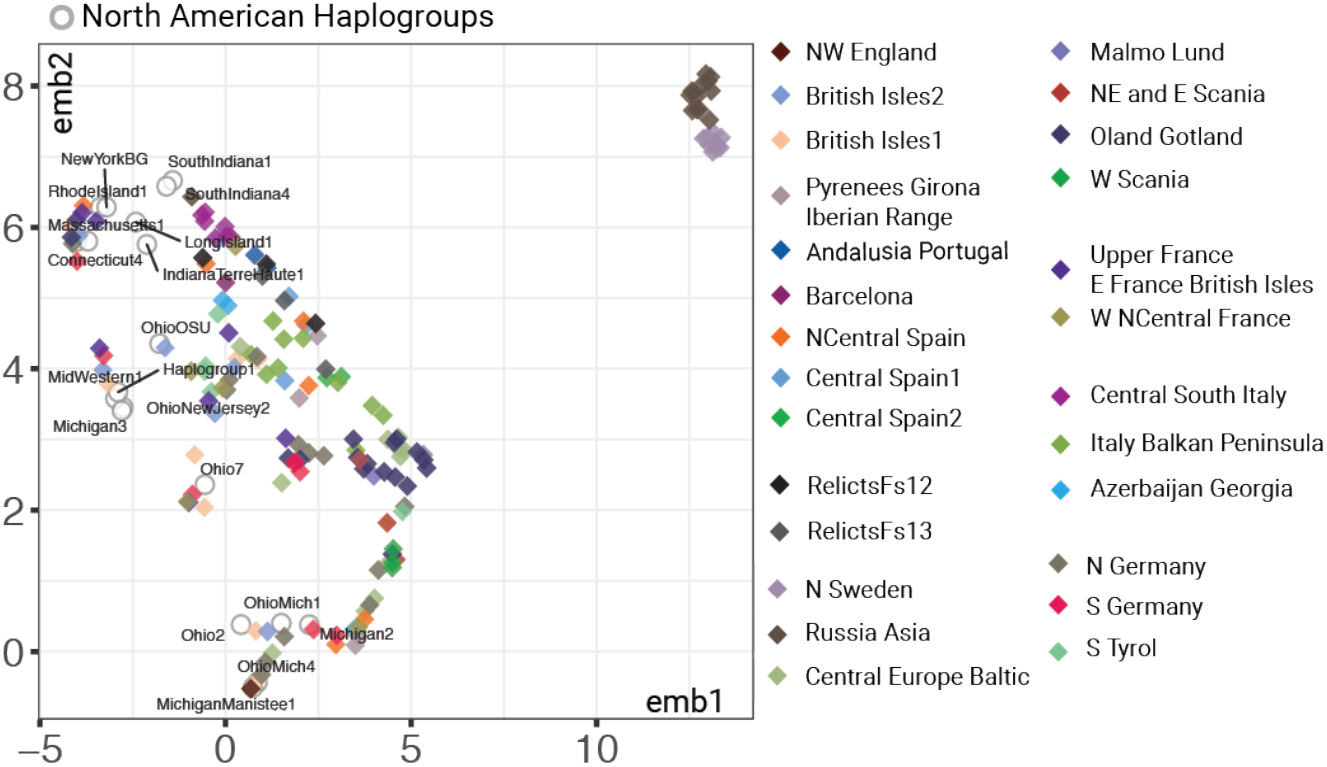
Projection of populations consisting of N. American groups and AEA *sub-clusters* in reduced environmental dimensions. UMAP embeddings showing populations in reduced dimensions of environmental variables T_avg_ (^°^C), precipitation (mm), and water vapor pressure (kPa).

### Current diversity has potentially reduced the burden of deleterious alleles

Evolutionary theory predicts that during range expansions and new colonizations deleterious mutations accumulate gradually and steadily, resulting in increased mutational load that can be reduced again by outcrossing (Peischl et al. 2013). We hypothesized that the levels of mutational load in N. American individuals would be related to rates of historic outcrossing as inferred from admixture. We estimated mutational load in each N. American *group* from the fraction of fixed derived deleterious alleles among all derived deleterious alleles. In addition, we looked at the relationship between this fraction and nucleotide diversity in each *group* (Fig. 5).

We observed a strong negative correlation (Kendall’s **τ** =−0.36) between the fraction of fixed among all deleterious derived alleles and nucleotide diversity. As expected, the *groups* that are sources of admixture, such as Hpg1, OhioMich1 and SouthIndiana4, had a relatively higher proportion of fixed derived deleterious alleles (Fig. 5, Fig. S12A) coupled with extensive reduction in nucleotide diversity. On the other hand the *groups* that emerged as a result of admixture among these and other lineages mostly showed higher nucleotide diversity and lower fixed derived deleterious to total derived deleterious ratio.

We specifically explored populations INRC and MISJ further, to look into the population frequency of derived deleterious alleles. Both populations are composed of multiple groups, with Hpg1, SouthIndiana4 and OhioMich1 as source lineages. Groups SouthIndiana2, IndianaNewJersey1 and SouthIndiana3 are admixture products of Hpg1 and SouthIndiana4, as indicated by *f*_*3* (Hpg1, SouthIndiana4: X)_ scores of −0.453, −0.411, −0.454 and *Z*-scores of −41, −30, −44. Similarly, groups OhioMich5 and OhioMich6 appear to be products of admixture between Hpg1 and OhioMich1 (*f*_*3* (Hpg1, OhioMich1: X)_ scores of −0.476 and −0.477, *Z-*scores of −138 and −128). In both populations, source lineages had higher proportions of derived deleterious alleles that are fixed in those groups (Fig. S12B,C), whereas these deleterious alleles were at intermediate frequencies in the admixed groups. This strongly suggests that admixture has relieved some of the potential mutational load that could have been caused by past fixation of deleterious alleles.

### Ongoing selection at several immunity loci in N. America

Apart from admixture reducing mutational load, it can also be a source of beneficial alleles. If such alleles are strongly selected, they will create signatures of a selective sweep (J. M. Smith and Haigh 1974; Stephan 2019; Moest et al. 2020). To look for such a scenario, we focused on large populations comprising several *groups* that apparently arose as a result of admixture between lineages that diverged before their introduction to N. America (Fig. S3). These populations were INRC (Indiana), NJSC (New Jersey), MISJ (Michigan), OHML and OHPR (both Ohio).

Methods that track the decay of haplotype homozygosity in a population (Vatsiou, Bazin, and Gaggiotti 2016) can be used to detect such sweeps. We scanned whole genomes for signals of natural selection using haplotype homozygosity based tests *iHS* (integrated haplotype homozygosity score) *(Voight et al. 2006)* and *nS*_*L*_ (number of segregating sites-by-length) (Ferrer-Admetlla et al. 2014) for individual populations (Table S5-9) and *xp-EHH* (cross population extended haplotype homozygosity) (Sabeti et al. 2007) for comparisons between population pairs. For individual populations, we focused on variants with |*iHS*| *p*-values for <0.001 and |*nS*_*L*_| values >2 (Table S5-S9). GO-term analysis of the 82 genes tagged by these variants revealed an enrichment of genes in the categories ‘response to stress’ and ‘response to stimulus’ (*p*-value after Bonferroni correction < 0.001 and FDR < 0.05) (Fig. 7A, and Table S10).

**Fig. 6.**
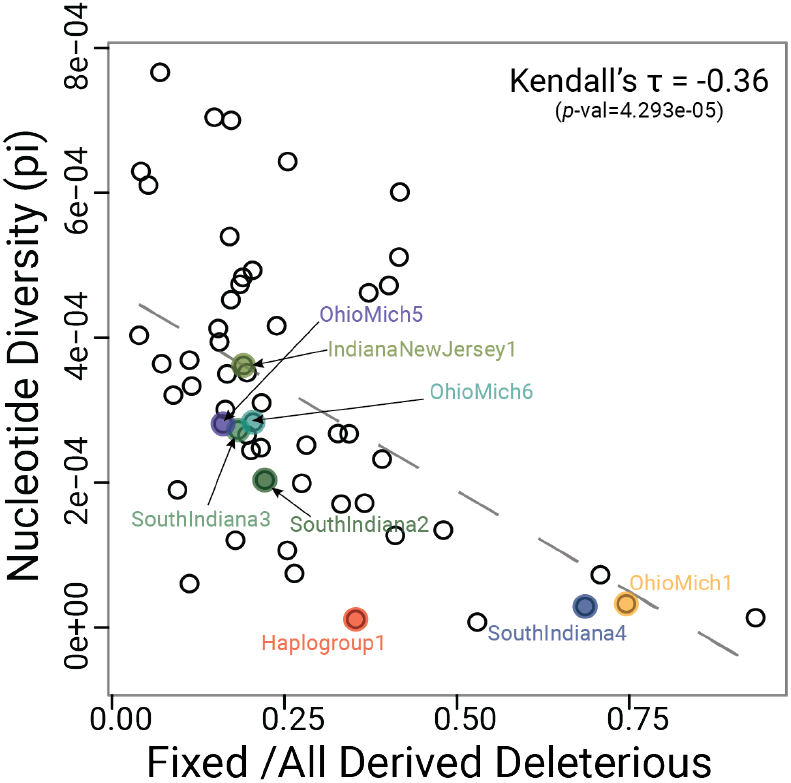
Newly formed groups have reduced mutational load and increased nucleotide diversity. Mean nucleotide diversity (pi) per group was calculated from estimation in 50 kb windows. Fixed derived deleterious (Fixed Der.Del.) corresponds to derived allele frequency = 1. Open circles represent each N. American group. Haplogroup1 and SouthIndiana4 are source groups of admixture in groups SouthIndiana3, SouthIndiana2 and IndianaNewJersey1. Haplogroup1 and OhioMich1 are source groups of admixture in groups OhioMich5 and OhioMich6.

**Fig. 7.**
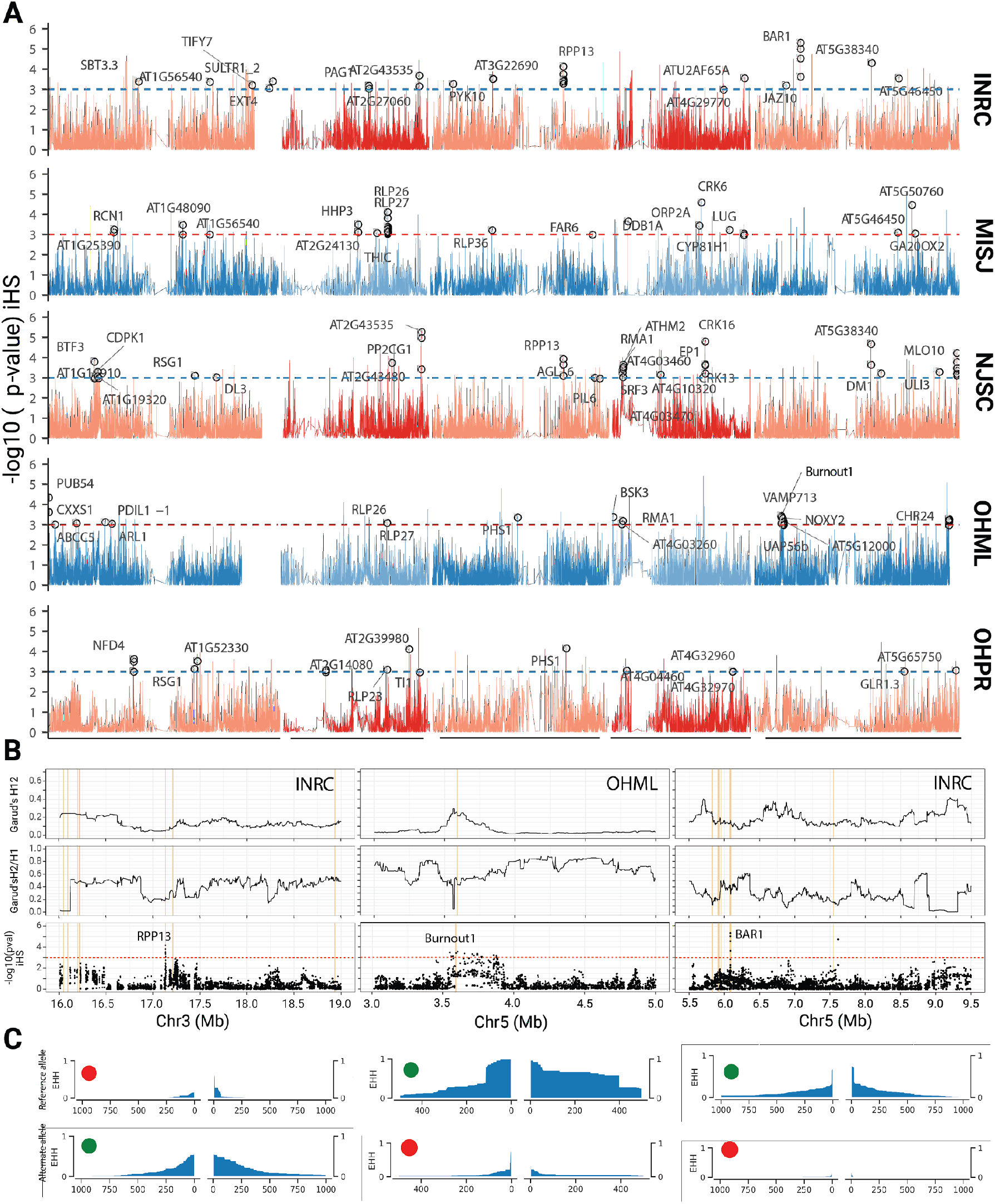
Genome-wide haplotype-based selection statistics in five N. American populations. **A.** Genome-wide *p*-values of |*iHS*| scores (based on empirical distribution). Dashed horizontal lines correspond to a *p*-value significance threshold of 0.001. Selection candidates, which also had |*n*S_L_| scores of >2, from the enriched GO categories of response to stress and response to stimulus (Fisher’s exact test with Bonferroni correction, *p*-value <0.001 and FDR <0.05) are plotted with gene names or gene IDs. **B**. Garud’s H_12_ and H_2_/H_1_ statistics and |iHS| scores around three disease resistance/NLR genes. **C**. Decay of extended haplotype homozygosity (EHH) around the variants with the lowest |iHS| *p*-value in the three genes shown in (B). Green dot indicates allele with selected variant, and red dot alternative allele. X-axis gives distance from focal variant as number of flanking variants in data set.

Consistent with the GO category enrichment, we noticed several NLR genes, a family that includes many known disease resistance genes (Van de Weyer et al. 2019). These included *RPP13* and *BAR1*, which confer resistance to the oomycete *Hyaloperonospora arabidopsidis* (Bittner-Eddy et al. 2000) and bacteria of the genus *Pseudomonas* (Laflamme et al. 2020). We measured the frequency of alternative haplotypes around these loci to determine the nature of the selective sweep (Garud et al. 2015). The most frequent haplotype is designated as H_1_ and the second most as H_2_, from which a modified product of haplotype frequency (H_12_) and the H_2_/H_1_ ratio are calculated. At *RPP13* and *BAR1*, we observed relatively low values for these two metrics and the presence of the selected alleles on multiple backgrounds, which together suggests soft sweeps at these loci (Fig. 7B,C). On the other hand, a pronounced hard sweep was observed in and around another putatively selected NLR, *BURNOUT1*, in the population OHML (Fig 7B,C), with the selected allele found on a single haplotype.

Similar to the |*iHS*| and |*nS*_*L*_| results, genes with high *xp-*EHH scores included several genes known to be involved in biotic and abiotic stress responses (Table S11).

## Discussion

How newly introduced, non-indigenous species adapt to new environments is a topic of long-standing interest in the field of eco-evolutionary biology of invasive species (Baker, H. G. & Stebbins, G. L. 1965; Bock et al. 2015). There are two potential challenges facing invasive species: First, the niches in the new environment might be different from the ones in the native range and/or already filled by other species. Second, introductions typically begin with few individuals and therefore potentially a narrow genetic basis. The initial lack of genetic diversity can be overcome by new mutations or through the generation of new genetic combinations, either by crosses among the introduced population or with close relatives that are present in and already adapted to the new environment. We have used *A. thaliana* to address these questions.

*Arabidopsis thaliana* is native to Europe, Asia and Africa, where it is found mostly as a human commensal (Hoffmann 2002; 1001 Genomes Consortium 2016; Lee et al. 2017; Durvasula et al. 2017; Zou et al. 2017). The human-assisted expansion of this species to N. America presents an excellent system to study processes associated with colonization of a new environment because it occurred recently and because the genetic diversity in the native range is so well documented for *A. thaliana*. Previous work has laid the groundwork for our study, but was limited by a paucity of genetic markers (Platt et al. 2010) or a focus on a single dominant lineage (Exposito-Alonso et al. 2018). We have investigated multiple individuals from several N. American populations at the whole-genome level, allowing us to describe fine-scale haplotype sharing within N. America and between N. America and individuals from the native range, either sequenced as a part of the 1001 Genomes project (1001 Genomes Consortium 2016) or subsequent efforts focused on Africa (Durvasula et al. 2017), China (Zou et al. 2017) and Ireland (this work).

### Multiple independent introductions

The extant diversity among *A. thaliana* individuals in N. America can be traced back to multiple, almost certainly independent introductions of lineages of diverged ancestries from three distinct geographic regions of Western Europe (British Isles/Ireland, Upper and Eastern France), central Europe (Germany, Czechia, and Austria) and Eastern Europe (the Baltic region and Russia). We detected these introductions using haplotype-based methods (Lawson et al. 2012), allele frequency-based methods (Patterson et al. 2012) and a method based on rare-allele sharing (Schiffels et al. 2016; Flegontov et al. 2019), lending considerable confidence to our findings and illuminating the extant diversity from several different angles. Significantly, even though we confirm that North-Western Europe and specifically the British isles are a major source of multiple introductions, the predominant lineage Hpg1, which has been estimated to have been introduced ~400 years ago (Exposito-Alonso et al. 2018), has more ancestry from Upper and Eastern France than from the British Isles. Our approach of haplotype-based clustering of individuals at different hierarchical levels using fineSTRUCTURE (Lawson et al. 2012) has allowed us to pinpoint several Western European sources of N. American *A. thaliana*. While the sparser representation of individuals from Eastern Europe and Asia has limited our ability to more precisely identify the source of introductions from these regions, it is clear that Eastern Europe has contributed to extant N. American *A. thaliana* ancestry. Historical patterns of human migration indicate that northern and western Europeans arrived in significant numbers from 1840s to 1880s followed by waves of southern and eastern Europeans from the 1880s to 1910s (Passel and Fix 1994), which are reflected in the genetic make-up of present-day humans in N. America (Bryc et al. 2015; Dai et al. 2020). In the regions where we collected *A. thaliana* in N. America humans have more British, Irish, central and eastern European ancestry than western, southern and northern European ancestry (Bryc et al. 2015), consistent with the *A. thaliana* ancestry patterns. Thus, local anthropogenic introduction of *A. thaliana* can be accepted as a parsimonious explanation for the presence of diverged lineages in the regions of N. America that we sampled in our study.

### Wide-spread admixture

Perhaps our most significant finding is how multiple introductions have led to present-day N. American *A. thaliana* being surprisingly genetically diverse, different from many other colonizing or invading species (K. M. Dlugosch and Parker 2008). This highlights how between-population variation in the native range has translated into within-population variation in N. America (Rius and Darling 2014). In organisms with low out-crossing rates such as *A. thaliana,* benefits of local adaptation in the native range hinder admixture from other populations, even in the face of inbreeding depression. It has been argued that during invasion of new territory, there is a temporary loss of local adaptation that not only lifts the maladaptive burden of admixture but even favors admixture (Verhoeven et al. 2011; Rius and Darling 2014). Similar patterns as we have reported here for *A. thaliana* have been suggested for other systems, mostly based on limited genetic information and without the benefit of being able to infer ancestry along each chromosome (B. Facon et al. 2008; Kolbe et al. 2004; Lavergne and Molofsky 2007; A. L. Smith et al. 2020).

Based on the observation of lower selfing rates in N. America compared to Europe, it has been suggested that under slightly increased outcrossing, mixing of haplotypes should be expected (Platt et al. 2010). In line with this hypothesis, we observed that most N. American *A. thaliana* populations have individuals with admixture from the dominant Hpg1 haplogroup. Being apparently already well-adapted to the N. American ecological context upon its introduction, today Hpg1 is wide-spread lineage in N. America (Platt et al. 2010; Exposito-Alonso et al. 2018). Admixture with Hpg1, followed by selection, might have benefitted and accelerated the spread of new incoming lineages. Alternative explanations such as short-term fitness benefits through heterosis (B. Facon et al. 2005; Keller and Taylor 2010) can currently not be ruled out, but could be tested with common garden experiments across N. American field sites.

### Purging of deleterious mutations

An important aspect of colonization is the severe genetic bottleneck due to founder effects and subsequent accumulation of deleterious mutations (Kirkpatrick and Jarne 2000; Schrieber and Lachmuth 2017; Verhoeven et al. 2011; Willi 2013), further exacerbated by predominant self-fertilization (Noël et al. 2017). One of the ways out of this invasion paradox (Estoup et al. 2016) might be admixture between colonizing lineages, which can both remove deleterious mutations (Heller and Maynard Smith 1978) and generate new genetic combinations that are only adaptive in the new environment (K. M. Dlugosch and Parker 2008; Rius and Darling 2014). Consistent with these predictions, we observed that the admixed N. American *A. thaliana* haplogroups have fewer fixed derived deleterious alleles and increased genetic diversity, as measured in terms of nucleotide diversity. This finding demonstrates that admixture has been successful in purging some of the potential mutational load carried by the founder lineages. A caveat is that deleteriousness of variants is based on presumed reduction or loss of molecular function (Kono et al. 2018), even though gene inactivation can be adaptive as well (Olson 1999; Weigel and Nordborg 2015). A more direct approach to determining the extent of purging of mutational load in N. American colonizing lineages could come from direct estimates of local adaptation deficits and selection coefficients, by comparing the fitness of N. American individuals at their site of collection against a global sample of *A. thaliana* accessions (Exposito-Alonso et al. 2019) or by quantifying the amount of genetic rescue or F_1_ heterosis in crosses between populations, see (Koski et al. 2019).

### Resistance genes as loci under selection

An indication of selection having potentially shaped the geographic distribution of genetic diversity in N. American *A. thaliana* is the observation of environmental dissimilarity between N. American haplogroups and their source lineages from the native range being negatively correlated with shared ancestry between them. Given that *A. thaliana* is a human commensal in its native range, it is not hard to envision that anthropogenically induced adaptation to invade (AIAI) (Hufbauer et al. 2012) might play a significant role in having accelerated *A. thaliana*’ adaptation to the N. American environment.

If a species is far from an adaptive peak, large-effect mutations are particularly likely to affect progress of adaptation (Fisher 1930). While the relative importance of abiotic and biotic factors for adaptation is still debated (Morris et al. 2020), some of the most drastic effects arise from disease resistance genes, where single genes have outsized effects on fitness and survival on plants in the presence of pathogens. In *Capsella*, it has been shown that dramatic losses of genetic diversity after extreme genetic bottlenecks can be tolerated at most genes in the genome, except for immunity loci (Koenig et al. 2019). Our selection scans with *A. thaliana* individuals from five different N. American populations have revealed that genes related to biotic stress are enriched among selection candidates. These include genes known to have alleles that confer resistance genes to two of the most prominent pathogens of *A. thaliana*, *H. arabidopsidis* and *Pseudomonas* (Holub and Beynon 1997; T. L. Karasov et al. 2014; Talia L. Karasov et al. 2018). One of the loci we found to be under selection is *RPP13* (Rose et al. 2004), whose product recognizes the co-evolved, highly polymorphic effector ATR13 from *H. arabidopsidis* (Allen et al. 2004). Another one is *BAR1*, whose product recognizes members from the conserved HopB effector family from *Pseudomonas* (Laflamme et al. 2020). While *RPP13* is under balancing selection in at least part of the native range (Allen et al. 2004), we observe that an *RPP13* allele is maintained on different haplotypes and has undergone a selective sweep in North American *A. thaliana* populations. Given that *H. arabidopsidis* appears to be an *A. thaliana* specialist (Slusarenko and Schlaich 2003), it must have been introduced with its *A. thaliana* host, and its genetic diversity in the introduced range might as low or even lower than that of its host, potentially providing an explanation for the apparent selective sweep at *RPP13*.

### Conclusions

Altogether, our analysis using whole-genome sequences from extant N. American *A. thaliana* has established a scenario of multiple introductions from sources of previously diverged Eurasian lineages. The distribution of diversity in N. America appears to mirror that of recent human migrants from Eurasia to N. America. We provide evidence that new haplotype diversity has been generated through wide-spread admixture among introduced lineages, relieving mutational load and providing raw material for selection to act upon. Our findings are thus consistent with early plant scientists who proposed that hybridization can lead to the introduction of adaptive variation via introgression or admixture (Anderson 1948, 1949; Stebbins 1959; Grant 1981). The advent of molecular analyses has confirmed the importance of hybridization for adaptation and speciation (M. L. Arnold 1996, 2004; Rieseberg 1997) and here we demonstrate the potential importance of admixture to invasive success. Admixture has been shown to facilitate successful colonization when individuals from divergent populations have been recurrently introduced to a new range (Rius and Darling 2014; Katrina M. Dlugosch et al. 2015; Estoup et al. 2016). North American *A. thaliana* therefore may not have suffered from the genetic paradox of invasion (Allendorf and Lundquist 2003; Estoup et al. 2016). *Arabidopsis thaliana* has also colonized other continents, including S. America and Australia (Kasulin et al. 2017; Alonso-Blanco and Koornneef 2000), and it will be interesting to determine both how genetic diversity of *A. thaliana* in these other places compares with N. America, and how genetic diversity of *A. thaliana* compares with that of other plants that have been inadvertently introduced to N. America by humans (La Sorte, Mckinney, and Pyšek 2007; Neuffer and Hurka 1999; Durka et al. 2005).

## Methods

### Sample collection and sequencing

Briefly, two strategies were used for the sample collections: 1. Herbarium collection: whole rosettes were collected from the locations and dried by pressing in acid-free paper with wooden press for 8-12 weeks. 2. Fresh tissue collection: 2-3 well expanded leaves from a field plant were collected in a microcentrifuge tube and frozen on dry ice (at the site) and later kept at −80°C until further processing. Seeds of Irish collection were a gift of Sureshkumar Balasubramanian (Monash University, Australia) were stratified in 0.1% agar for 7 days and grown on soil for 3 weeks at 23°C in short days (8 hours light, 16 hours dark). A fully expanded leaf was used for DNA extraction from these plants. Genomic DNA from plants collected in herbarium sheets was done following PTB extraction method (Kistler 2012). Genomic DNA from 2-3 mg fresh tissue was isolated with the protocol described previously (Clarke 2009).

#### Sequencing

Libraries for reduced-representation (RAD-seq) with KpnI enzyme were prepared by following the protocol (Rowan et al. 2017) with a modification of 100ng as starting concentration of DNA instead of 200ng to target sequencing depth to 25-30x. Whole-genome sequencing (WGS) was performed with average targeted coverage of 12-15x using protocol adapted from (Talia L. Karasov et al. 2018). Libraries were sequenced on Illumina HiSeq2000 (RAD-seq: 1 x 100 bp and 1 x 150bp single-end) and HiSeq3000 (WGS: 2 x 150bp paired-end) platforms. Summary of sequencing in terms of mean depth per individual is in Table S1. Additionally, short reads (fastq) for selected *A. thaliana* from Africa (Durvasula et al. 2017), China (Zou et al. 2017), herbarium (Exposito-Alonso et al. 2018) and outgroup *A. lyrata* (Novikova et al. 2016) genomes were downloaded from NCBI-Sequence Read Archive using sratoolkit.2.8.2. Short reads of 145 accessions, a subset of the 1,135 genomes of the 1001 Genome collection (1001 Genomes Consortium 2016) were downloaded from the servers in the Max Planck Institute for Developmental Biology.

### Mapping and variant calling

Single-read (generated by sequencing RAD-seq libraries) and paired-end read DNA sequences (generated by sequencing WGS libraries in this work and downloaded from NCBI-SRA) in fastq format were mapped using bwa-mem (bwa-0.7.15) algorithm (Heng Li 2009) to TAIR10 reference genome (https://www.arabidopsis.org/download_files/Genes/TAIR10_genome_release) and sorted using samtools v1.3 (Li et al. 2009). Illumina paired-end reads from herbarium accessions were additionally trimmed with skewer (v. 0.1.127) (Jiang et al. 2014) using default parameters and merged with flash (v. 1.2.11) (Magoč and Salzberg 2011) with a maximum overlapping value of 150 bp, prior to the mapping step as described before. SNP calling was performed following Genome Analysis ToolKit (GATK) best practices with slight modifications to accommodate for single-read sequencing (DePristo et al. 2011; Van der Auwera et al. 2013). GATKv3.5 pipeline was used on individual samples with the following suite of tools: RealignerTargetCreator, IndelRealigner, HaplotypeCaller, SelectVariants, VariantFiltration, BaseRecalibrator, PrintReads. Individual gvcf files thus generated were used for joint genotyping using GenotypeGVCFs tool of GATKv3.5 resulting in a merged variant call format (vcf) file that stored all the variant calls. Detailed parameters used during the SNP calling and filtering are described in the script provided in the accompanying repository.

#### Inclusion of SNPs from remaining A. thaliana global diversity dataset

Once the initial set of SNPs was called from the samples collected and sequenced in this project together with African, Chinese, Irish, 1,135 subset and outgroup *A. lyrata*, additional SNP filtering was performed. This filtering procedure retained SNPs that are: 1. Bi-allelic, 2. missing less than 70% data, 3. Not from genomic regions annotated as transposable elements (TAIR10 annotations) and 2000 bp up and down-stream from these annotations 4. QD (Quality-by-depth) greater than 20. After this filtering 1,159,256 SNPs were retained. Further, to include the remaining 990 individuals from 1,135 genomes, SNP calls from publicly available SNP dataset (1001 Genomes Consortium 2016) were lifted from the 1,159,256 positions. This introduced some sites to be multi-allelic, therefore, additional filtering was performed using vcftools v0.1.15 (Danecek et al. 2011) to retain SNPs that are only bi-allelic and additionally those SNPs which had more than 10% missing data were discarded. The filtering strategy employed here resulted in very high-quality 862,934 SNPs with average genotyping rate of 0.94 in total of 1689 individuals.

#### Estimation of recombination rates

Haplotype phasing for estimation of recombination rate was performed with ShapeIt2 (v2.r837) (Delaneau et al. 2013). Samples from this project and a subset of 1135 genomes (1001 Genomes Consortium 2016) samples for which raw reads were processed in the same pipeline as described in this project, were used for the analysis. A slightly different SNP and individual filtering strategy was used. First, only individuals with average sequencing coverage (depth) >10x were kept. Then, the ollowing SNP filters were applied: 1. No singleton positions. 2. No SNPs from TAIR10 annotated transposable elements (TE) and SNPs 2,000 bp up and down-stream of annotated TEs. 3. No SNPs from NBS-LRR (NLR) clusters with mean depth above 25. The additional filtering at individual and variant level was performed to avoid over-estimation of switch error rate (incorrectly phased heterozygous sites) because of the sensitivity of Shapeit2 to coverage and quality (Delaneau et al. 2013). After phasing, the recombination rate variation along the chromosomes was estimated using LDhelmet v1.7. Mutation matrix for LDhelmet analysis was calculated using a parsimony-based method (Chan, Jenkins, and Song 2012). In brief, if two plant genotypes of *Arabidopsis lyrata* (outgroup) shared the same allele, it was assigned as ancestral and then these sites were used to calculate the mutation probability matrix. LDhelmet output is population scale recombination rate (*ρ* = 4*Ne.r*) in 1/bp units. For downstream analysis, we scaled this recombination rate in cM/Mb units by applying frequency-weighted means method (Booker, Ness, and Keightley 2017) on an empirical recombination map of *A. thaliana* F_2_ mapping populations (Salomé et al. 2012). The recombination map is available in the accompanying repository.

### Population genetic analysis

#### PCA, UMAP and IBD

Principal Component Analysis (PCA) was performed using SmartPCA of EIGENSOFT version 6.0.1 (Patterson, Price, and Reich 2006) package. SmartPCA was used with and without 2 outlier iterations. Outlier iterations were performed to remove some of the highly diverged individuals from Iberian and North African relict populations. For UMAP analysis on the same dataset, we first performed PCA using Python package sklearn v 0.23.2 (Pedregosa et al. 2011). Then, we used the first fifty PCs as input for generating two UMAP embeddings using Python package umap v0.4.6 (McInnes, Healy, and Melville 2018). Details of the analyses are in the supplementary methods. Identity-by-descent and identity-by-state analysis was carried out with PLINK v1.90 (Chang et al. 2015).

#### Chromosome painting and clustering

Clustering of individuals based on shared ancestry from haplotype data was performed using fineSTRUCTURE on a coancestry matrix derived with the software CHROMOPAINTER v2 (Lawson et al. 2012). Reconstruction of each individual’s ancestry based on haplotype sharing is summarized by CHROMOPAINTER v2 in a coancestry matrix. CHROMOPAINTER v2 treats all the individuals (except the individual whose ancestry is being reconstructed) as donor haplotypes and generates a mosaic of shared chunks copied from these donors in a given recipient individual. Similarity in the patterns of shared chunks (copying vectors) is indicative of shared ancestry and is the basis of the model-based clustering approach taken by the fineSTRUCTURE algorithm. Specifically, we performed this analysis in the following hierarchical way:

1. All the N. American individual haplotypes were painted as a mosaic of all the other N. American individuals’ haplotypes (self-excluding)
2. All the non-N. American (Afro-Eur-Asian /AEA) haplotypes were formed as a mosaic of each other. Based on the haplotype sharing these individuals were then clustered and grouped into what we call *sub-clusters*, *clusters* (comprised of *sub-clusters*) and *regions* (comprised of clusters representing specific geographical regions)
3. All the N. American haplotypes were then formed as a mosaic of AEA haplotype clusters. Detailed description of the analysis is in supplementary methods.

#### Treemix analysis

Treemix infers the relationship among populations as a graph structure derived from genome-wide allele frequency and genetic drift modeled as Gaussian distribution (Pickrell and Pritchard 2012). We determined the phylogenetic relationship among the N. American *groups* as inferred by fineSTRUCTURE algorithm using Treemix v1.13. After processing the dataset with helper scripts available in the repository https://bitbucket.org/nygcresearch/treemix/wiki/Home, we used *A. lyrata* individuals as outgroup and set blocks of 100 SNPs to calculate the maximum likelihood tree using 6 bootstraps.

#### *f*_*3*_-outgroup analysis

In order to determine the extent of shared drift between the Afro-Eur-Asian (AEA) sub-clusters (smallest fineSTRUCTURE grouping) and N. American haplogroups, we used *f*_*3*_-outgroup tests as described (Patterson et al. 2012). *N. American _(i)_, AEA SubCluster _(j)_: Relicts (Fs12_3)* configuration was used and implementation of the test was carried out using R package “*admixr”* (Petr, Vernot, and Kelso 2019).

#### qpWave and D-statistic analysis

In order to determine minimum number of ancestry waves from Afro-Eur-Asian (AEA) regions (comprised of different haplogroup *sub-clusters* defined by fineSTRUCTURE analyses) we used *D*-statistic and qpWave analysis from ADMIXTOOLS software (Reich et al. 2012). The analyses were performed in a hierarchical way. First a set of AEA regions were chosen as outgroups (outgroup_layer1) that were donating the haplotype chunks at equal proportions to the analyzed N. American haplogroups. All the N. American haplogroup pairs were analyzed with this set of outgroups (*N. American1, N. American2:Outgroup1, Outgroup2*). Composition of this outgroup set is described in the Table S4. The population pairs for which Rank 0 matrix was accepted were considered to be from a single stream of ancestry compared to the outgroups (*p*-value >0.05). These pairs were then further analyzed one-by-one in another round of qpWave analysis with the same outgroup set plus one more region with higher contribution of haplotype chunks across the North American haplogroups (Fig. 3). These additional regions were: 1. BritishIsles2; 2. Italy/Balkan Peninsula; 3. NorthGermany; 4. RussiaAsia; 5. Upper/EastFrance BritishIsles.

#### Rare allele sharing

1039 AEA individuals that formed the fineSTRUCTURE *sub-clusters* were used as a reference panel to ascertain rare alleles and calculate rare allele sharing (RAS) between AEA *sub-clusters* and N. American haplogroups. The input files were prepared with the tools from repository at (https://github.com/stschiff/rarecoal-tools) and the analysis was performed by the pipeline available at (https://github.com/TCLamnidis/RAStools). Minimum allele count of 2 and maximum allele count of 20 was used on the SNPs with less than 10 % missing data. Alleles were polarized with the *A. lyrata* data.

### Environmental factor analysis

Historical climate data from 1970-2000 were downloaded from WorldClim2.0 (Fick and Hijmans 2017) at 2.5-minute resolution using Python library latlon-utils 0.0.5 (https://github.com/Chilipp/latlon-utils). Environmental variables average temperature (°C), precipitation (mm), solar radiation (kJ m^−2^ day^−1^) and water vapor pressure (kPa) were used for further analysis. Pairwise Euclidean distances of all the environmental variables were calculated for each N. American haplogroup to AEA *sub-clusters* (mean Latitude −Longitude of individuals in a given *sub-cluster* was used) and standardized values were used to model shared drift (measured by outgroup-*f*_*3*_ statistics) among N. American haplogroups and AEA *sub-clusters* as a function of the environmental variables using Bayesian multilevel (hierarchical) linear regression. Description of the priors and hyper-priors is in the supplementary methods.

Projection of the N. American haplogroups in reduced dimension formed by standardized average temperature, precipitation and vapor pressure was performed using uniform manifold approximation and projection (UMAP) (McInnes, Healy, and Melville 2018). Two independent runs of UMAP were performed with different random numbers. In both the runs default "Euclidean" distance was used to compute distances in high dimensional space. Details of the scripts and notebooks used for the analysis are in the accompanying repository.

### Estimation of derived deleterious allele frequency estimation

To determine ancestral state of the positions, pairwise alignments between *A. thaliana* (TAIR10) and *A. halleri* and between *A. thaliana* (TAIR10) and *A. lyrata* were obtained from (ftp://ftp.ensemblgenomes.org/pub/plants/release-44/maf/ensembl-compara/pairwise_alignments/). These alignments were then converted to BAM format using maf-convert tool (Kiełbasa et al. 2011) and samtools (Li et al. 2009) was used to convert the alignment in BAM format to a VCF file which carried variant information. If both the *A. halleri* and *A. lyrata* had the same homozygous reference allele (with respect to TAIR10) at a given position, then that allele was considered ancestral. In case of alternate alleles, either the homozygous alternate alleles in both *A. halleri* and *A. lyrata* were considered as ancestral, or, if there were homozygous alternate alleles in one and heterozygous ones in the other, the consensus was considered to be ancestral (scripts and ancestral states are in the accompanying repository). Precomputed SIFT 4G predictions for effect of mutations for *A. thaliana* (TAIR10) were obtained from (https://sift.bii.a-star.edu.sg/sift4g/public//Arabidopsis_thaliana/). Using these predictions, positions with deleterious effect were selected and their derived allele frequency in each N. American haplogroups was calculated. Nucleotide diversity in these haplogroups was independently calculated in 50 kb windows using vcftools v1.15 (Danecek et al. 2011).

### Genome-wide selection scans

We performed haplotype homozygosity based selection scans to detect recent and ongoing selection. *iHS* (integrated haplotype score) (Voight et al. 2006) and XP-EHH (cross-population extended haplotype homozygosity) (Sabeti et al. 2007) were calculated using hapbin (Maclean, Chue Hong, and Prendergast 2015), details are described in the supplementary methods. Recombination map generated earlier was used in the estimation of both the statistics. *nS*_*L*_ (number of segregating sites by length) (Ferrer-Admetlla et al. 2014), Garud’s H1, H12 and H2/H1 (Garud et al. 2015) (window size = 500, step size= 10), Tajima’s D (window size=50000 and step size=5000) were calculated with scikit-allel (Miles et al. 2020). Nucleotide diversity for the population was calculated using a pipeline described by (Martin, Davey, and Jiggins 2015).

|*iHS*| and |*nS*_*L*_| were used in a complementary manner. As *iHS* is known to be affected by recombination rate variation (O’Reilly, Birney, and Balding 2008), we used *iHS* first and based on empirical distribution of the scores, *p*-values were calculated per SNP. *nS*_*L*_ was then calculated on the same dataset. As *nS*_*L*_ is robust to variation in mutation and recombination rates (Ferrer-Admetlla et al. 2014), overlap of the SNPs that showed |*iHS*| *p*-value less than 0.001 and|*nS*_*L*_| higher than 2 was taken as a signal of selection. GO-term analysis of the genes carrying the candidate selected SNPs was performed with AgriGOv2 (Tian et al. 2017) with PlantGo-Slim categories. For enrichment of GO terms Fisher’s exact test with Bonferroni correction was used.

## Data and Code Availability

Code and high resolution images from the main text are available from https://github.com/weigelworld/north_american_A.thaliana repository. Short reads have been deposited in the European Nucleotide Archive under the accession number PRJEB42417.

## Author Contributions

GS, JD and DW conceived the project. GS and JD organized the collection trips. GS, JD and CF collected the samples. GS, JD, AB, MQD, AGH and DL processed the samples for sequencing and performed DNA extractions. GS, JD and AB prepared sequencing libraries. GS and JD scripted and ran sequence processing pipelines. GS performed formal analysis. HB, CF and DW supervised the research. GS wrote the original draft. GS, SML, DSL, CF, HB and DW reviewed and edited the draft.

## Acknowledgements

We thank Rebecca Schwab for the help with the organization for the first collection trip; Claudia Friedemann (MPI for Developmental Biology), Cathy Herring (North Carolina State University Central Crops Research Station), Carol Ann McCormick (University of North Carolina), Jon Peter (New York Botanical Garden), Nelson Garcia (Rutgers University), Rosa Raudales and Sanjukta Majumder (University of Connecticut), Robert Capers (George Safford Torey Herbarium), Irina Kadis and Alexei Zinovjev (Arnold Arboretum), Jason Parrish, Jiangbo Fan, Guo-liang Wang, Lynn Jin, David Mackey, Maria Bellizzi and John Freudenstein (Ohio State University), Neville Millar (Kellogg Biological Station), Eric Knox (Indiana University), Timothy Morton and Joy Bergelson (University of Chicago), Alan Fryday, Bethany Huot and Sheng Yang He (Michigan State University), and Christine Niezgoda (Field Museum of Natural History) for help during sample collection; Suresh Balasubramanian (Monash University) for seeds of Irish accessions; Julian Regalado, Dino Jolic, Danelle Seymour, Jörg Hagmann and Jorge Quintana for help with setting up the RAD-seq analysis pipeline and for help throughout the project; Christa Lanz, Julia Hildebrandt for help with sequencing; and Talia Karasov and Richard Neher for discussions. This work was funded by the Max Planck Society.

## Supplementary Material

### Supplementary Methods

#### 1 Principal Component Analysis(PCA) and uniform manifold approximation and projection(UMAP)

##### 1.1 PCA

We used smartpca v.13050 for performing PCA on the merged RAD-seq and WGS datasets. Extremely diverged individuals of Relicts ancestry [1] affected the PCA strongly (PC1 and PC2 capturing differences between these individuals and the rest) and detailed population structure within the rest of the AEA individuals was not apparent in both the RAD-seq dataset (Fig. S2) and WGS. To overcome this, we used outlier removal procedure (Option: numoutlieriter=2) with default parameters of outlier sigma threshold of 6.0 and number of PC components to perform outlier iterations equal to 10. For projection of N. American individuals into the PC space formed by the AEA individuals, we also used above mentioned outlier removal procedure. We used individuals grouped into AEA ***clusters*** for the projection analysis and specified the list of these clusters in “poplistname” option in the smartpca parameter set-up file (this list is in the repository).

##### 1.2 UMAP

For UMAP analysis we processed the genotype vcf file with the scikit-allel package [2] of Python. First, we performed LD-based pruning with 500 SNP sliding window size, 50 SNP as step size and *r*^2^ threshold of 0.5, that resulted in retaining 200,000 SNPs after 5 iterations. PCA was performed on this dataset using package scikit-learn v 0.23.2 [3]. We used first 50 PC components for constructing two-dimensional embedding with number of neighbors =100 and minimum distance of 0.8 using Python package umap and umap-plot [7] (details in the notebook umapWGS.ipynb in the repository).

#### 2 Chromosome painting and clustering

##### 2.1 Generation of co-ancestry matrix with CHROMOPAINTER

###### 2.1.1 Estimation of *N*_*e*_ and *θ* parameters

We ran CHROMOPAINTERv2 to estimate the nuisance parameter *N*_*e*_ and *θ* using the expectation-maximization option (100 steps). Runs for all 5 chromosomes were performed independently. Final *N*_*e*_ = 9838.359 and *θ*= 0.01126963 values were calculated by weight-averaging (across each chromosome by size in *cM*) as described in [4]. Example command:

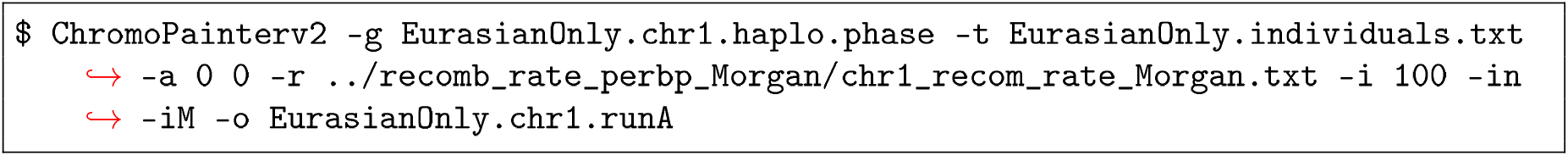

**Table 1:** *N*_*e*_ and *θ* parameters for all the chromosomes

| Chromosome | N_SNPs | $N_e$ | $\theta$ |
| --- | --- | --- | --- |
| 1 | 140484 | 11302.50 | 0.0110791 |
| 2 | 71243 | 6279.04 | 0.0111337 |
| 3 | 94479 | 14940.90 | 0.0116722 |
| 4 | 86040 | 6446.52 | 0.0120141 |
| 5 | 119015 | 8642.20 | 0.0107181 |

##### 2.1.2 Building co-ancestry matrix

We generated two separate co-ancestry matrices using CHROMOPAINTER v2 runs on each chromosome separately for every Afro-EurAsian (AEA) individual (in AEA subset) and for every North American individual (in N. American set) using other individuals as donors in the respective subsets for all the five chromosomes independently.Example command:

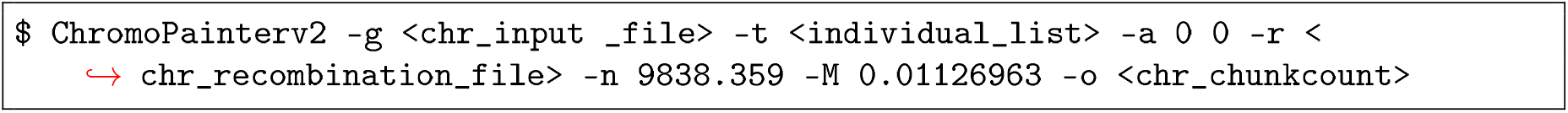

Co-ancestry matrices of five chromosomes were combined into one matrix to be used for clustering, using following command:

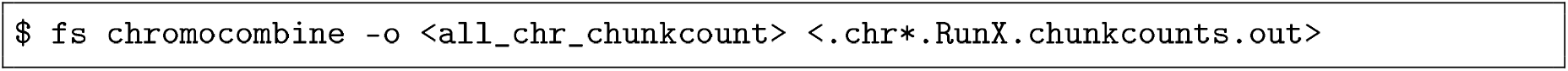

##### 2.2 Clustering with fineSTRUCTURE

###### 2.2.1 Identification of clusters from the individuals

We used output of co-ancestry matrix for further model-based Bayesian clustering of individuals in the groups using fineSTRUCTURE [5]. One million iterations were run with ten thousand iterations as burn-in and sampling was performed every ten thousand iterations. We performed 3 such independent runs by setting different seed (-s option) generated by random numbers. Example command:

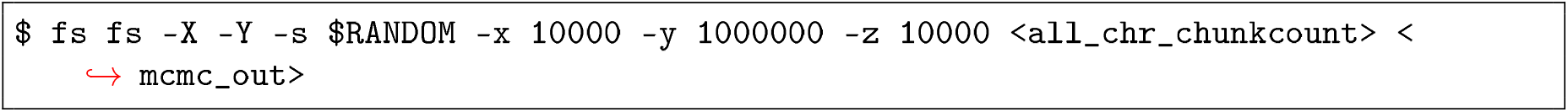

Following the clustering we produced a tree that summarized the relationships between the individuals classified in different clusters. We considered upto ten million trees for comparisons for splitting or merging individuals from the clusters. Again, three separate runs on previously generated outputs from three independent mcmc runs for clustering were performed. Example command:

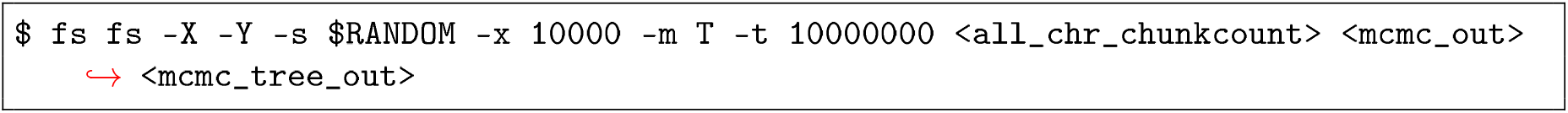

After each run final MAP states and mean coincidence for the tree file was generated using following example commands:

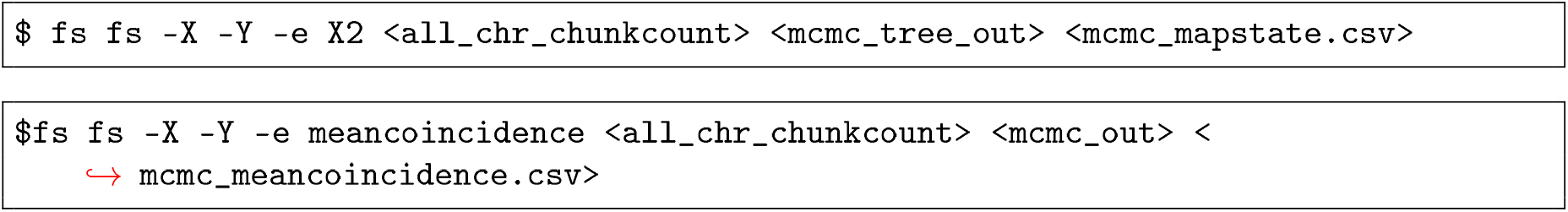

###### 2.2.2 Hierarchical ordering of clusters

We grouped individuals from AEA subset into 158 clusters that we call *sub-clusters* by visually inspecting the trees and merged these *sub-clusters* into 21 major clades that we call *clusters.* We further split these 21 major clades into 27 *regions* based on the geographical origin of the individuals in the *clusters.* The membership of individuals in these different *regions* is described in the Table S3. Similar strategy of visually inspecting tree was used to group N. American individuals into 58 clusters that we call *groups* (described in the Table S3). The *groups* were not further merged into clades.

##### 2.3 Chromosome painting of N. American *groups* with AEA *sub-clusters*

We applied CHROMOPAINTERv2’s ability to infer haplotype sharing among individuals to estimate N. American individuals’ copying profiles from AEA individuals, independently. We used following strategy:

i. We set each N. American *group* (comprised of member individuals) as a recipient and specified every individual from AEA-set as a donor according to its membership to *sub-cluster.* Thus, we specifically looked at haplotype chunks donated by AEA *sub-clusters* to N. American *groups* as a summary of haplotype segments donated by AEA individuals to N. American individuals. Following example command was used to accomplish this:

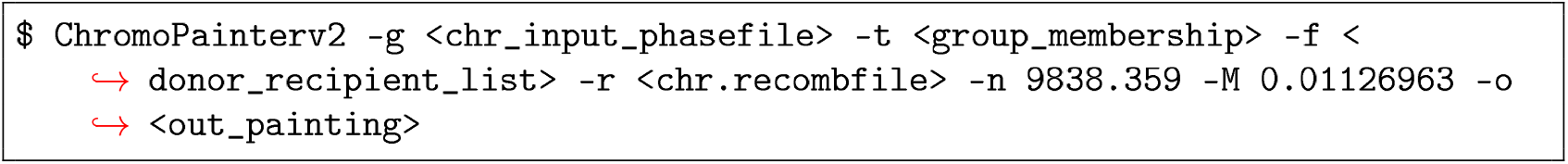
ii. After estimating counts of haplotype segments (chunks) shared by a N. American group with every AEA *sub-clusters,* we normalized this count by the number of individuals that formed a given AEA *sub-cluster.* These calculations were performed using custom python and bash scripts. (Uploaded in the repository)
iii. We calculated mean haplotype chunk counts donated by each AEA *region* by averaging over the haplotype segments donated by member *sub-clusters* of the *regions.* These calculations were performed using custom python and bash scripts. (Uploaded in the repository)

#### 3 Environmental factor analysis

##### 3.1 Bayesian multi-level linear regression model (outgroup *f*_*3*_ statistics as a function of environmental variables)

We modeled outgroup *f*_*3*_ as a Student’s T-distributed variable for *i*-th N. American group with normality parameter *v*_*i*_, location parameter *μ*_*i*_ and a global scale parameter *σ*

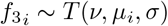

*μ*_*i*_ is expressed as linear combination of standardized environmental factor dissimilarity of the *i*-th N. American group with all the 158 AEA *sub-clusters* with corresponding *β* coefficients.

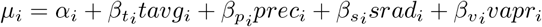

###### 3.1.1 Hyper-priors

We set following population wide hyper-priors.

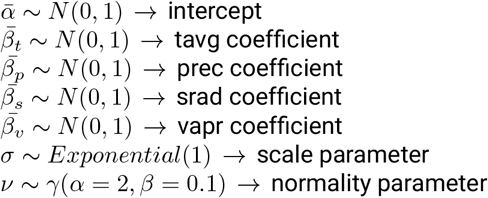

###### 3.1.2 Priors

We specified all the group-specific priors as normally distributed

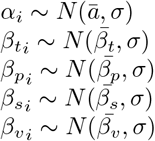

###### 3.1.3 Implementation

Implementation of the model was carried out in python using probabilistic programming package PyMC3 (v 3.9.3) [6]. For the analysis we sampled 4 chains with 1000 iterations for tuning and 4000 iterations as draws. Jupyter notebook (outGroupF3_hierarchicalRegression.ipynb) and related python scripts used to import and format the data are available in the accompanying repository.

##### 3.2 Projection of groups in reduced environmental space using UMAP

Bioclim data on the variables tavg, prec and vapr for each accession was downloaded according to its geographic location. We calculated mean of each variable for every individual and then according to the individuals’ membership to either AEA *sub-cluster* or N. American *group,* we grouped the individuals and calculated average for that *sub-cluster* or *group.* The projection of the *sub-clusters* and *groups* in reduced environmental space was performed with the python package umap-learn v0.4.6 [7]. umap_fs_newGroup_envVariableSpace.ipynb notebook in the repository describes the implementation.

#### 4 Genome-wide selection scans

We applied haplotype homozygosity based statistics to scan the genomes of individuals from select N. American populations (INRC, MISJ, NJSC, OHML and OHPR). *iHS* (integrated haplotype score) [8] calculates extended haplotype homozygosity (EHH) among given genomes while taking into account local recombination rates. Standardized *iHS* scores can be calculated by considering SNPs with similar allele frequencies [9]. We calculated *iHS* for every population with hapbin [10] where we set minimum allele frequency of 0.01 and EHH cutoff of 0.1 with other default parameters. Example command is:

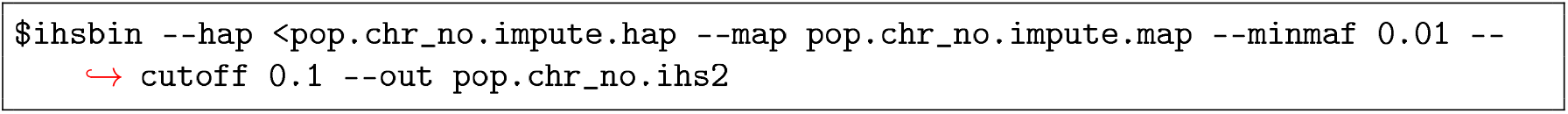

To complement the results from *iHS* we used standardized *nS*_*L*_ (number of segregating sites) [11] as implemented in Python package scikit-allel [2]. ***nS***_L_ is conceptually similar to *iHS* but instead of relying on local recombination rates, it relies on the number of adjacent polymorphic sites shared by a pair of haplotypes around focal SNP Garud’s H1,H12 and H2/H1 (in 500 SNPs window with step size of 10 SNPs) were calculated with the same package (nsl_GarudH_HaplotypeDiv_final.ipynb in the accompanying repository)

As selected allele approaches fixation, it becomes harder for *iHS* and *nS*_*L*_ to detect the signal of selective sweep. If the selected allele is fixed in one population and not in the other,then between-population comparison still can identify the signature of selective sweeps [9]. Therefore, we applied cross-population extended haplotype homozygosity (XP-EHH) test implemented in hapbin with default parameters. Example command:

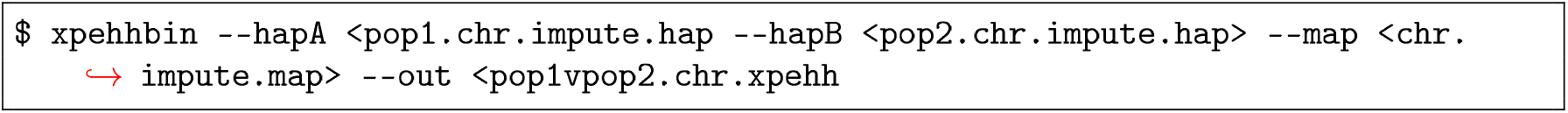

## List of Supplementary Tables

**Table S1.** Provenance and sequencing depth of individuals

**TableS2.** Scores of f3-statistic to test admixture among N. American *groups* in the form *(groupA,groupB:testGroup)* where z-scores are less than −3.0

**Table S3.** List of AEA sub-clusters and N. American *groups* and their member individuals

**Table S4.** Results of *qp-Wave* analysis

**Table S5.** |*iHS*| and |*nS*_*L*_| scores for SNPs showing significant |*iHS*| values in the population **NJSC** and gene IS of the SNPs

**Table S6.** |*iHS*| and |*nS*_*L*_| scores for SNPs showing significant |*iHS*| values in the population **MISJ** and gene IS of the SNPs

**Table S7.** |*iHS*| and |*nS*_*L*_| scores for SNPs showing significant |*iHS*| values in the population **OHPR** and gene ID of the SNPs

**Table S8.** |*iHS*| and |*nS*_*L*_| scores for SNPs showing significant |*iHS*| values in the population **OHML** and gene ID of the SNPs

**Table S9.** |*iHS*| and |*nS*_*L*_| scores scores for SNPs showing significant |*iHS*| values in the population **INRC** and gene ID of the SNPs

**Table S10.** GO term enrichment analysis for |iHS| significant SNPs (p-value less than 0.001)

**TableS11.** *xpEHH* values of SNPs and their *p*-values in cross-population comparisons along with the IDs of genes carrying these SNPs and their GO-description

Download of supplementary tables: https://nextcloud.tuebingen.mpg.de/index.php/s/Qy8PEL4Pkd6kibs

## List of Supplementary Figures

**Figure S1.**
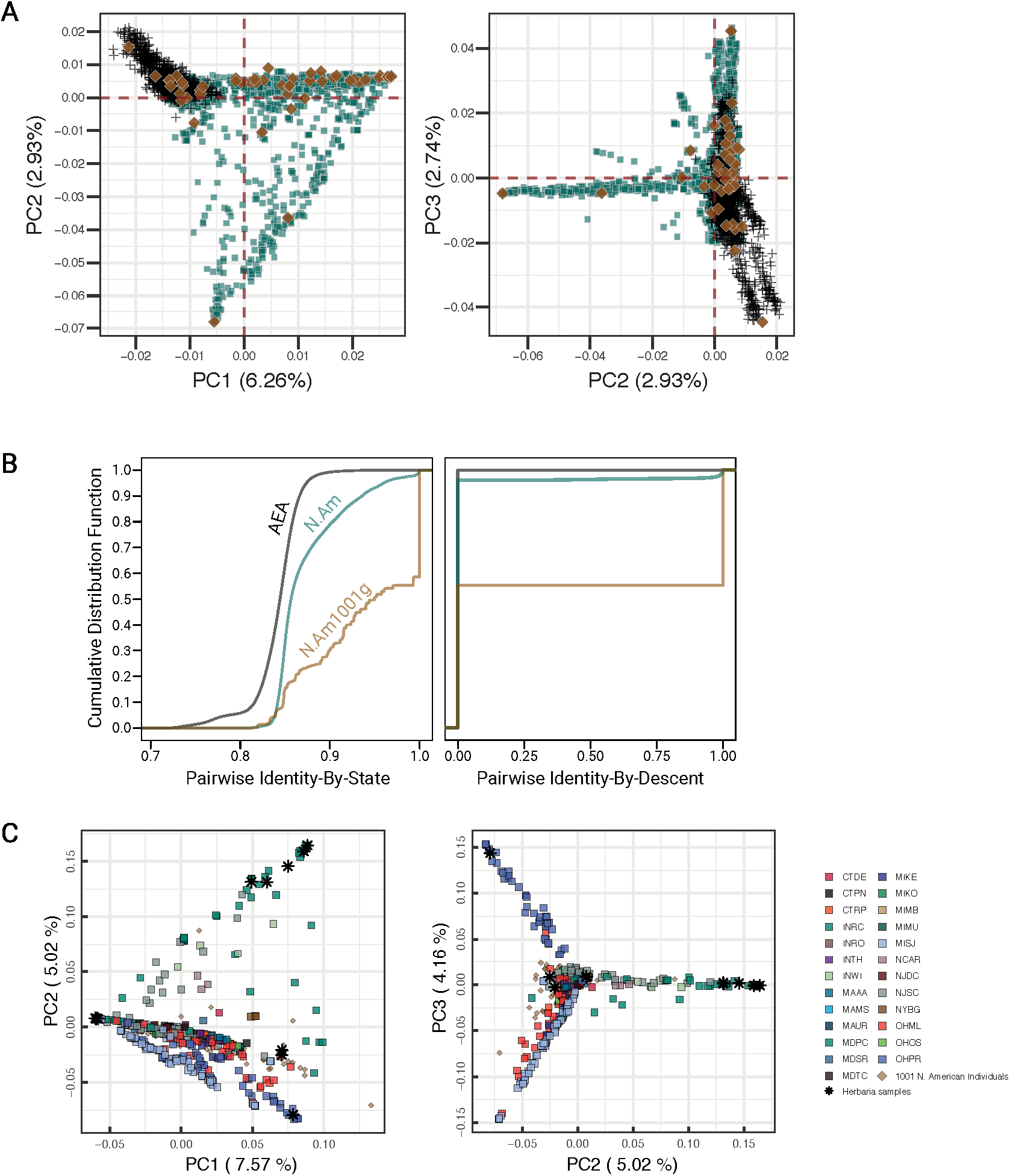
Population structure and genetic similarity of N. American populations in the context of global populations. **A.** Principal component analysis (PCA) with ~5,000 RAD-seq SNP markers with five outlier iterations in Eigenstrat SmartPCA. The individuals collected during this work in N. America are in blue (n = 3232). Brown: N. American individuals from 1001 Genomes (1001G) collection (n =135). Black: *Arabidopsis thaliana* from Afro-Eurasia (AEA, n=1,194). **B.** Empirical cumulative density function of genetic similarity estimated using pairwise Identity-by-State (DST) and Identity-by-descent among N American (blue) individuals, N. American (1001G individuals) and AEA individuals (black). **C.** PCA using whole genome sequences of 500 individuals (sampled in this work plus 1001G N. American individuals plus herbaria individuals) with ~900,000 SNP markers.

**Figure S2.**
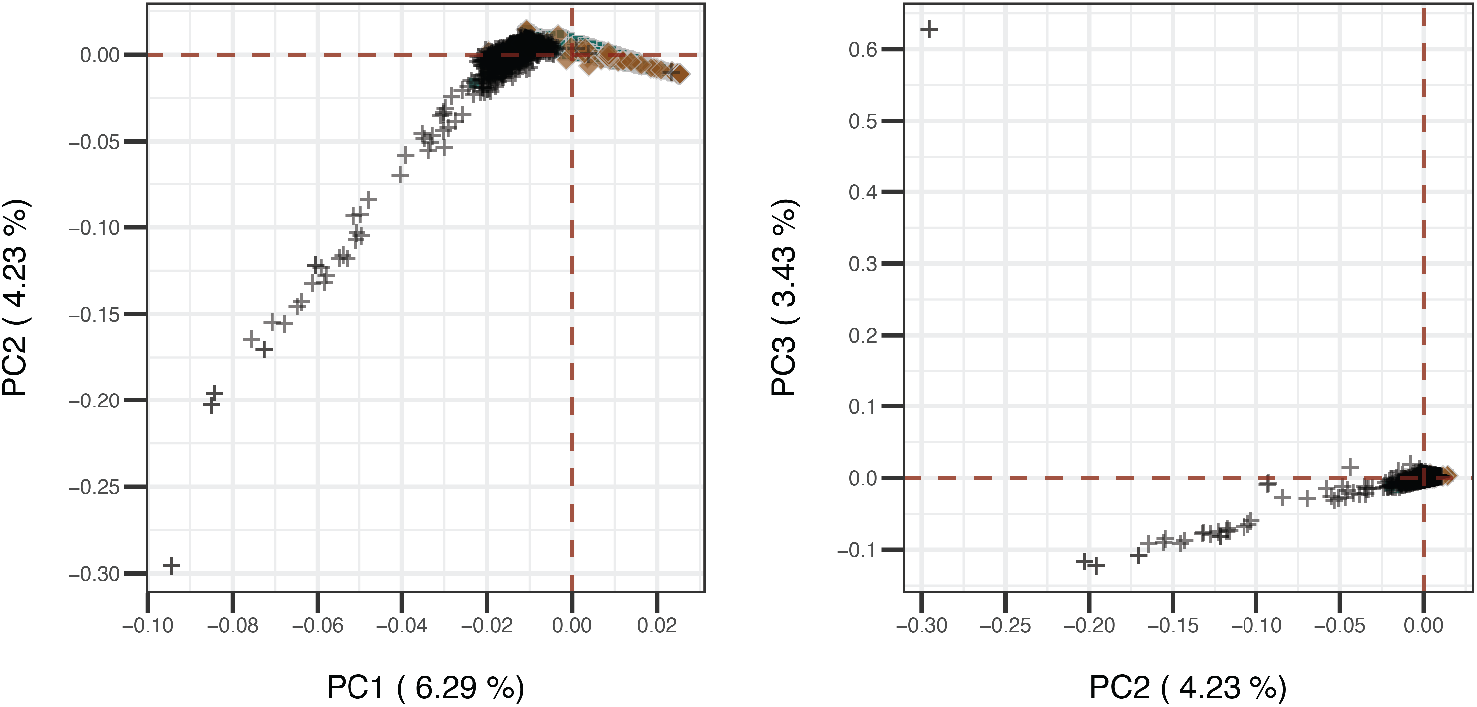
PCA using RAD-seq SNP markers with no outlier iterations implemented in SmartPCA. Blue, individuals collected in this work; brown: N. American individuals from 1001 Genomes collection (n=135); black: Afro-Eur-Asian individuals (n=1,194). Outliers detected here are individuals from relict populations from Iberian Peninsula, Sicily and sub-Saharan Africa.

**Figure S3.**
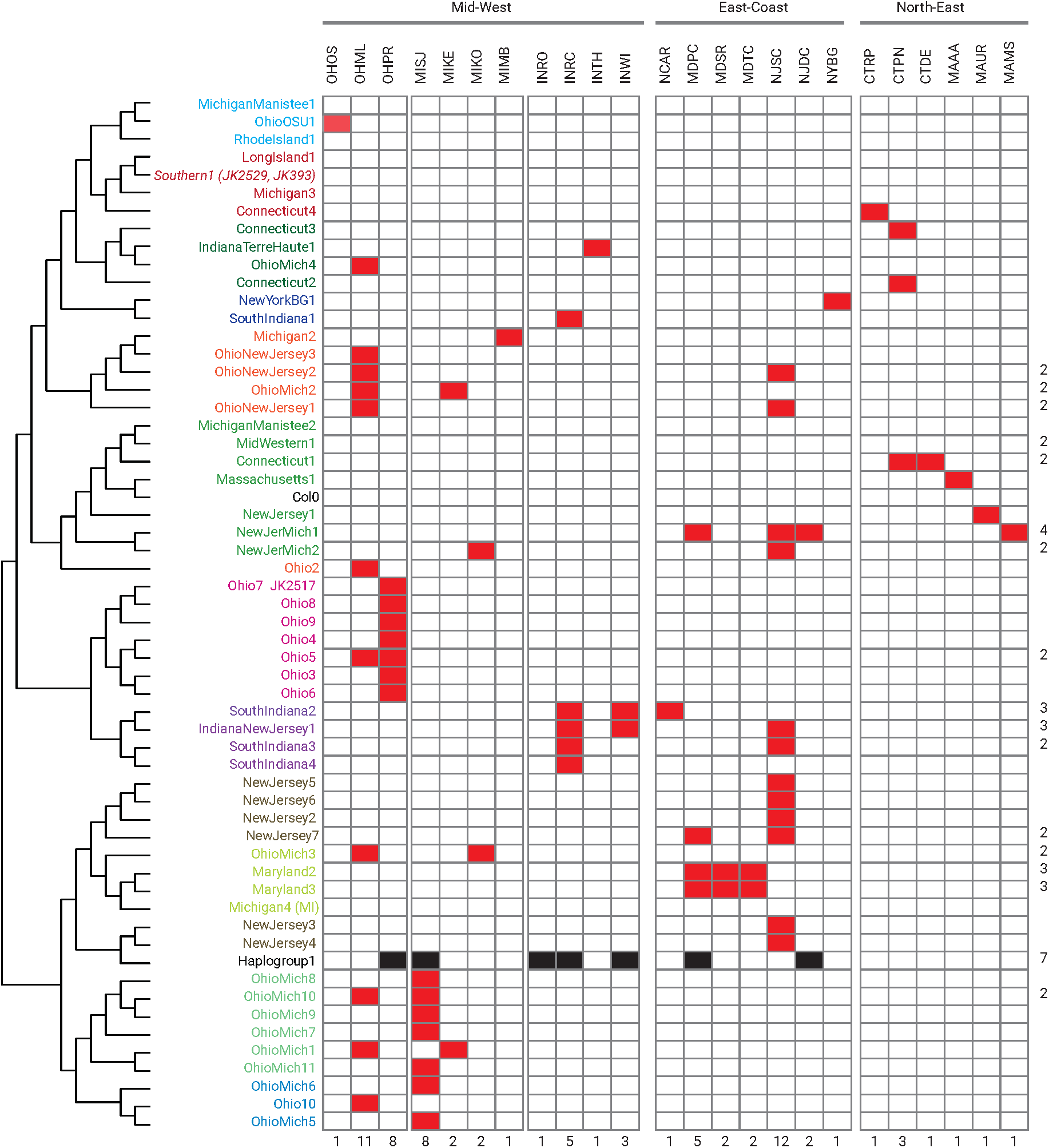
Summary of groups and populations. Last row of numbers represents the total count of *groups* present in the population and last column of numbers represents the number of populations in which a specific *group* is present (here a *group* present in a single population is not counted, count=1).

**Figure S4.**
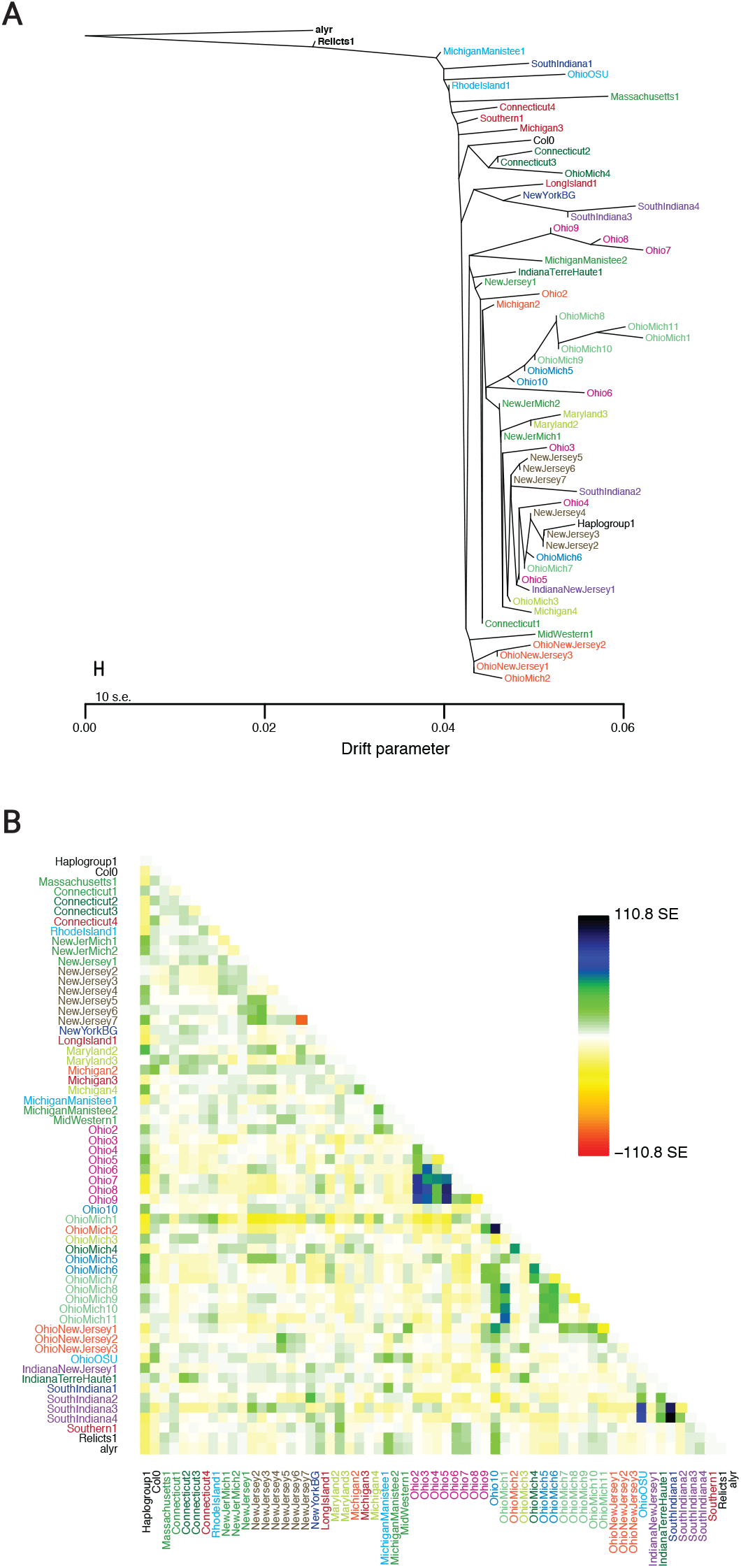
Maximum likelihood tree of N. American groups using Treemix Algorithm. **A.** Maximum likelihood (ML) tree of N. American groups defined by FINESTRUCTURE with no migration edges inferred, and *A. lyrata (alyr)* as outgroup. **B.** Residuals of ML tree. Residual covariance between any pair of groups derived from FINESTRUCTURE clustering divided by average standard error (in pairs, henca scaled residuals). Positive residuals indicate over-estimated covariance between a pair of populations (in green to dark blue shades), which are candidates for admixture.

**Figure S5.**
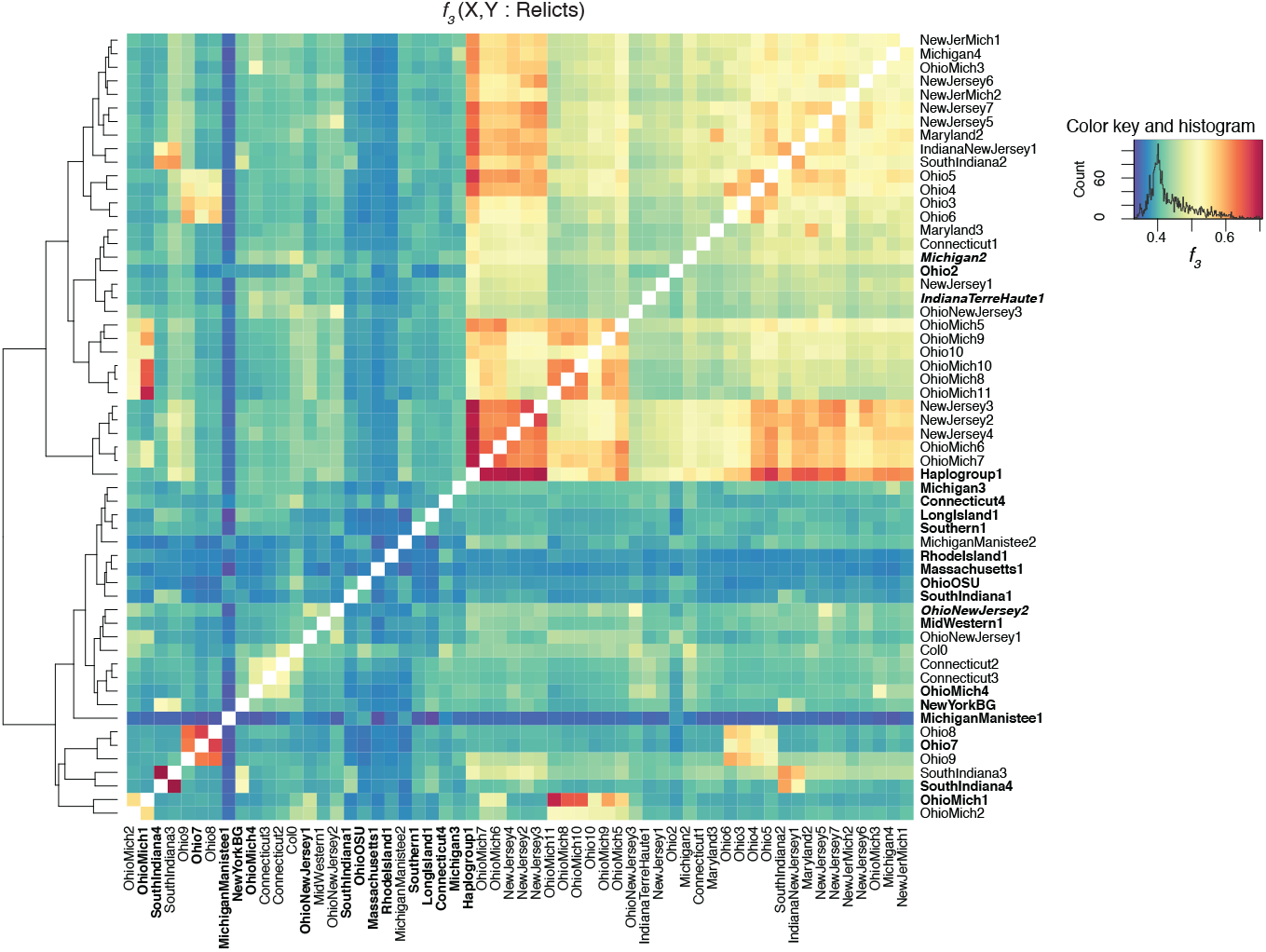
Shared drift among N. American haplogroups using *f*_*3*_ outgroup analysis. Outgroup *f*_*3*_ statistic in the form of (X,Y: relicts), where X and Y are two test N. American haplogroups. The heatmap shows the extent of shared drift among all the pairs of haplogroups (red: high shared drift; blue: low shared drift).

**Figure S6.**
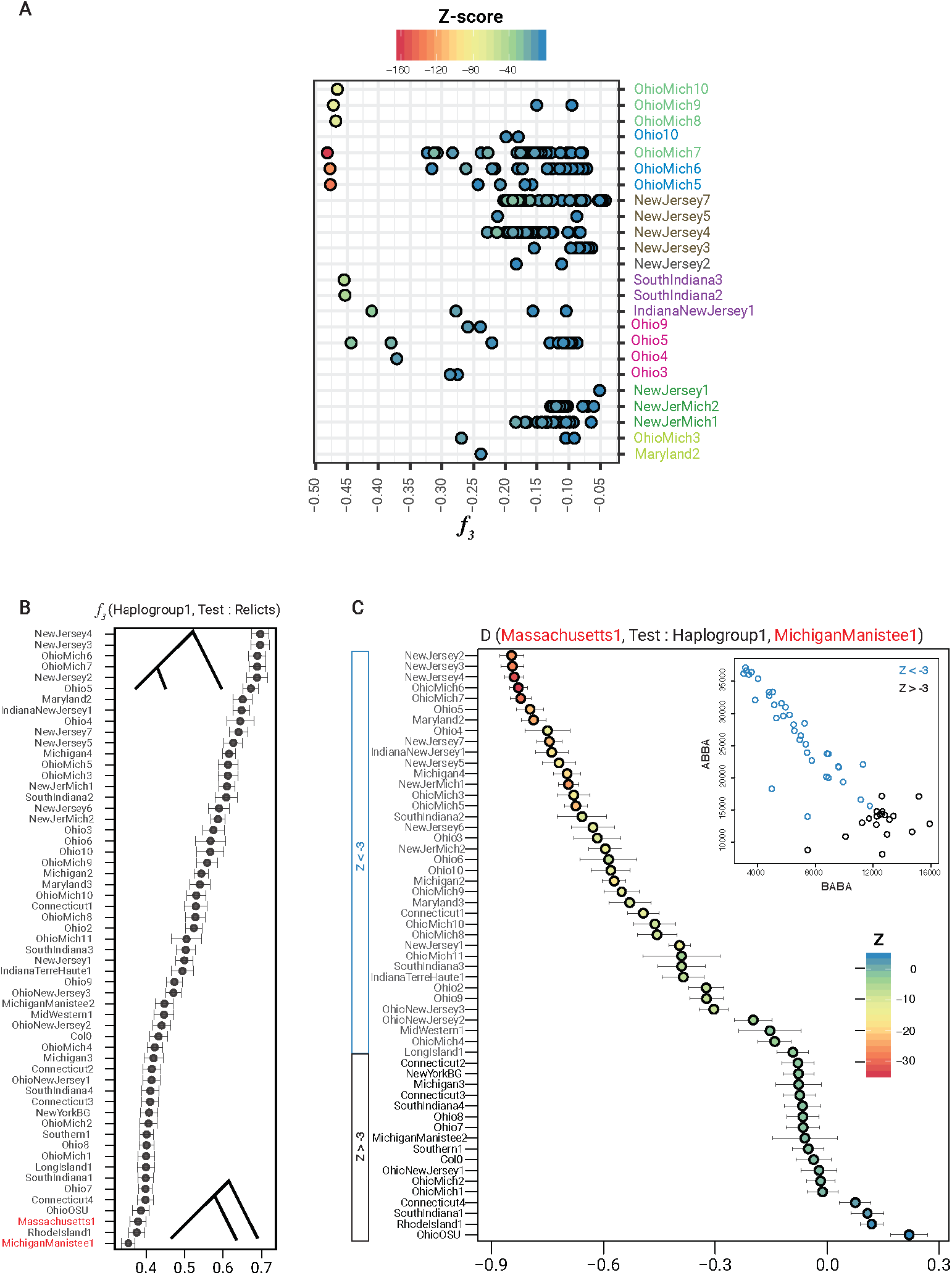
Admixed groups and shared drift of Haplogroup1 with other haplogroups. **A.** *f*_*3*_-statistic for detection of admixture in groups. Tests were performed with all the *f*_*3*_(*group*_*i*_,*group*_*j*_*: group*_*k*_) conlgurations possible where groups *i, j* and *k* are three distinct N. American groups. Scores of the groups (*group*_*k*_) with significant negative *f*_*3*_ scores (Z-score < −3) are shown, with Haplogroup1 as either *group*_*i*_ or *group*_*j*_ in the test configuration. **B.** Outgroup *f*_*3*_-statistic in the form of (Haplogroup1, test: relicts) to determine allele sharing between Haplogroup1 and other groups. **C.** D-statistics was then used in the form of (Massachusetts1, test: Haplogroup1, MichiganManistee1) to determine significant allele sharing (gene flow) between Haplogroup1 and test groups. Groups with Z-score < −3 are colored in blue and the rest in black. Inset: Count of BABA sites plotted against count of ABBA sites.

**Figure S7.**
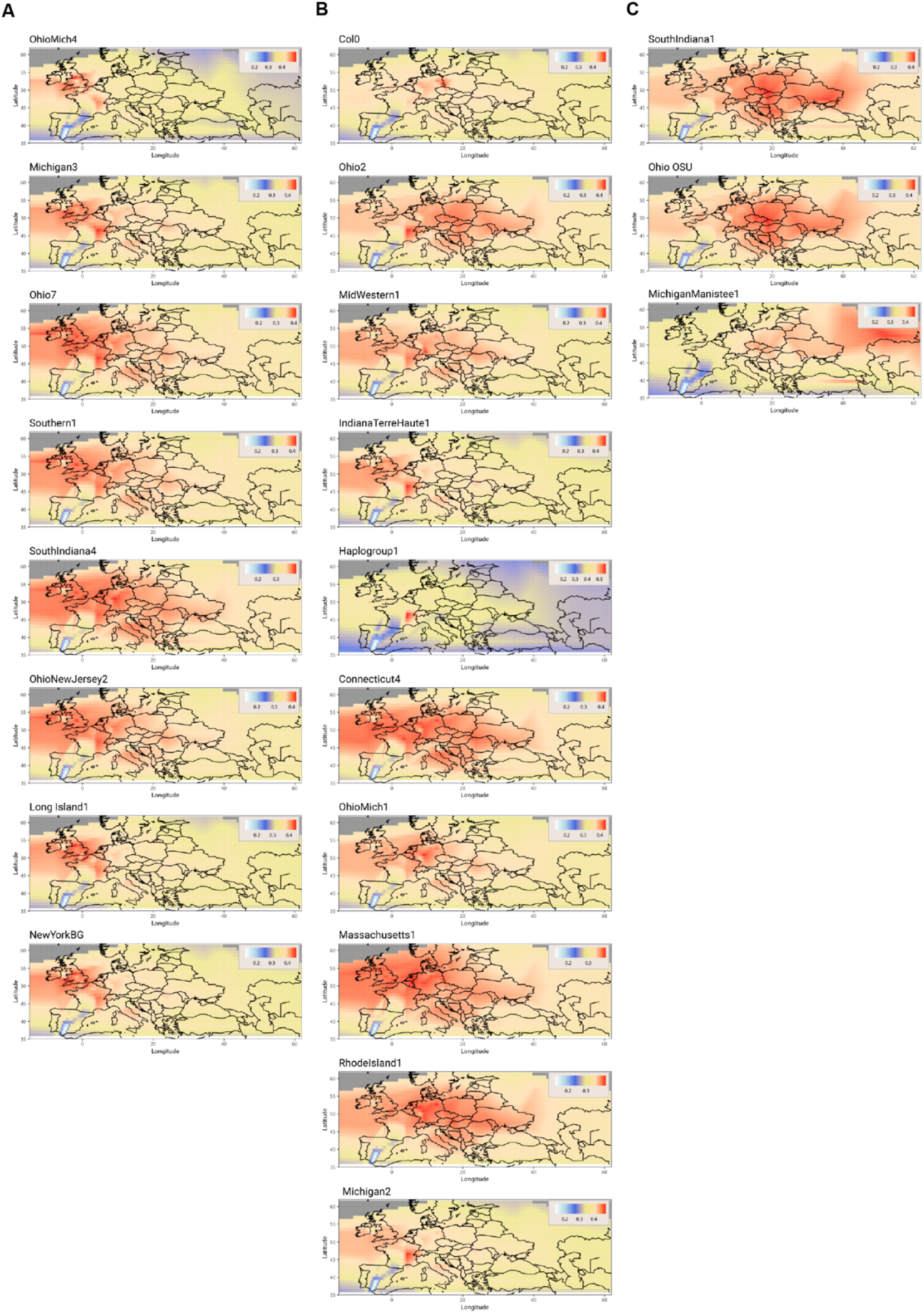
Shared drift of N. American haplogroups with AEA sub-clusters with *f*_*3*_ outgroup analysis. N. American haplogroups with excess shared drift (as inferred from *f*_*3*_-statistics) to **A.** Western Europe (mainly British Isles), **B.** central Europe, and **C.** Eastern Europe. Legends in the upper right corners show *f*_*3*_ values.

**Figure S8.**
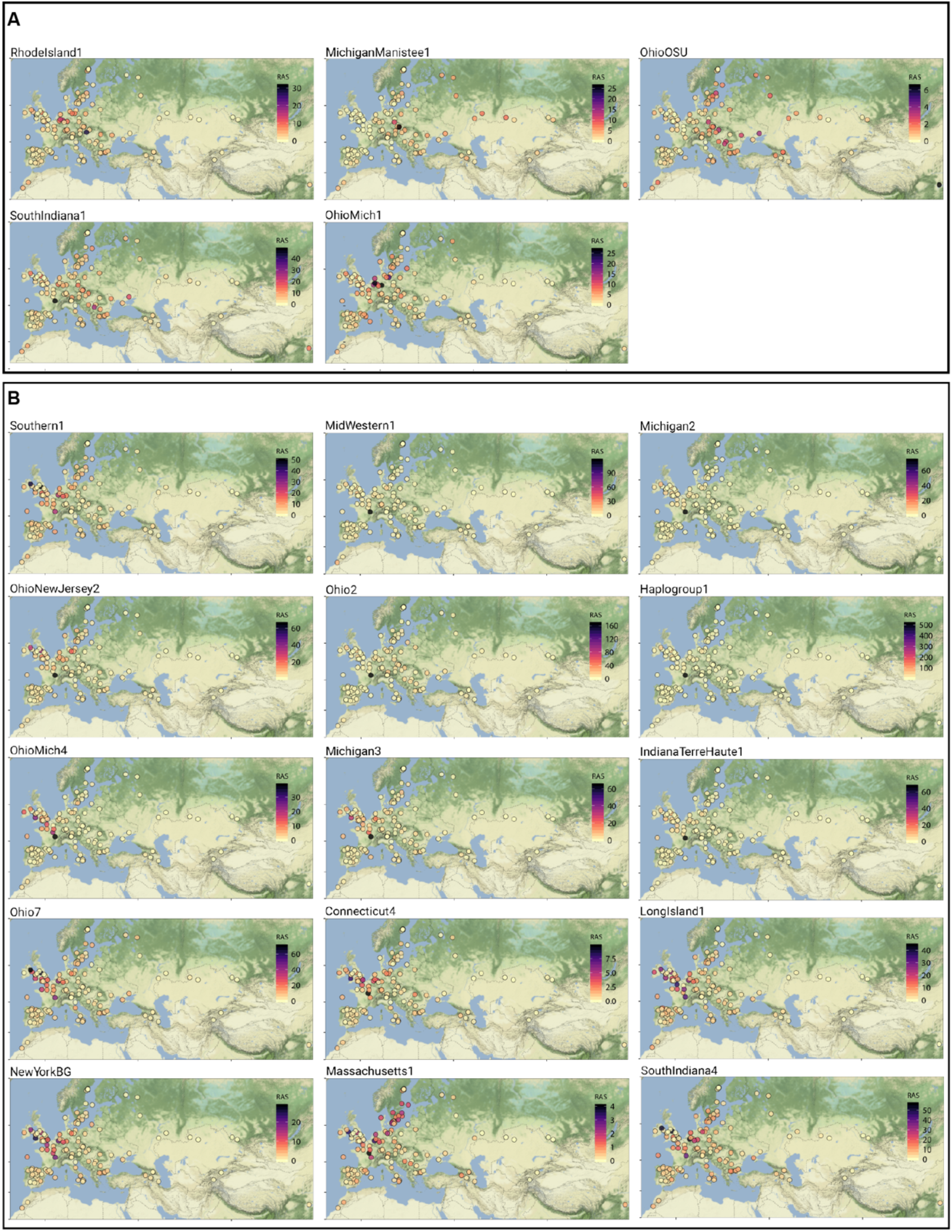
Rare allele sharing of N. American haplogroups with AEA sub-clusters. N. American haplogroups with excess rare allele sharing (RAS) to **A.** central/Eastern Europe, and **B.** Western Europe (mainly British Isles).

**Figure S9.**
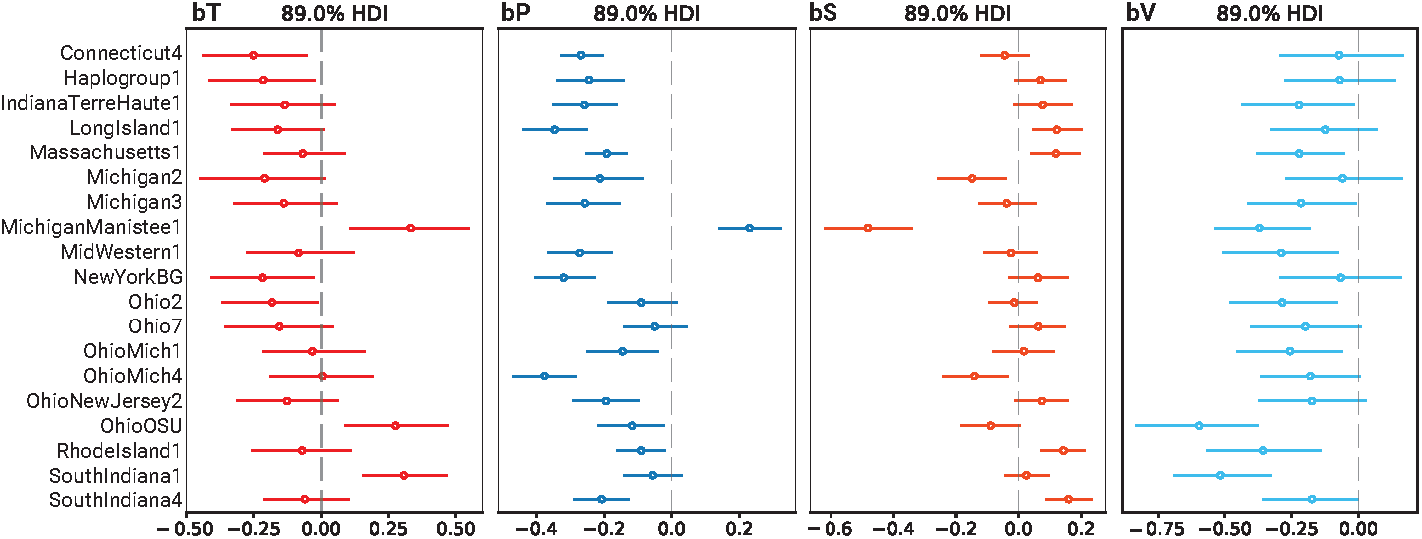
Posterior means and 89% compatibility interval for individual group’s *β* coefficients for tavg (bT), precipitation (bP), solar radiation (bS) and water vapor pressure (bV)

**Figure S10.**
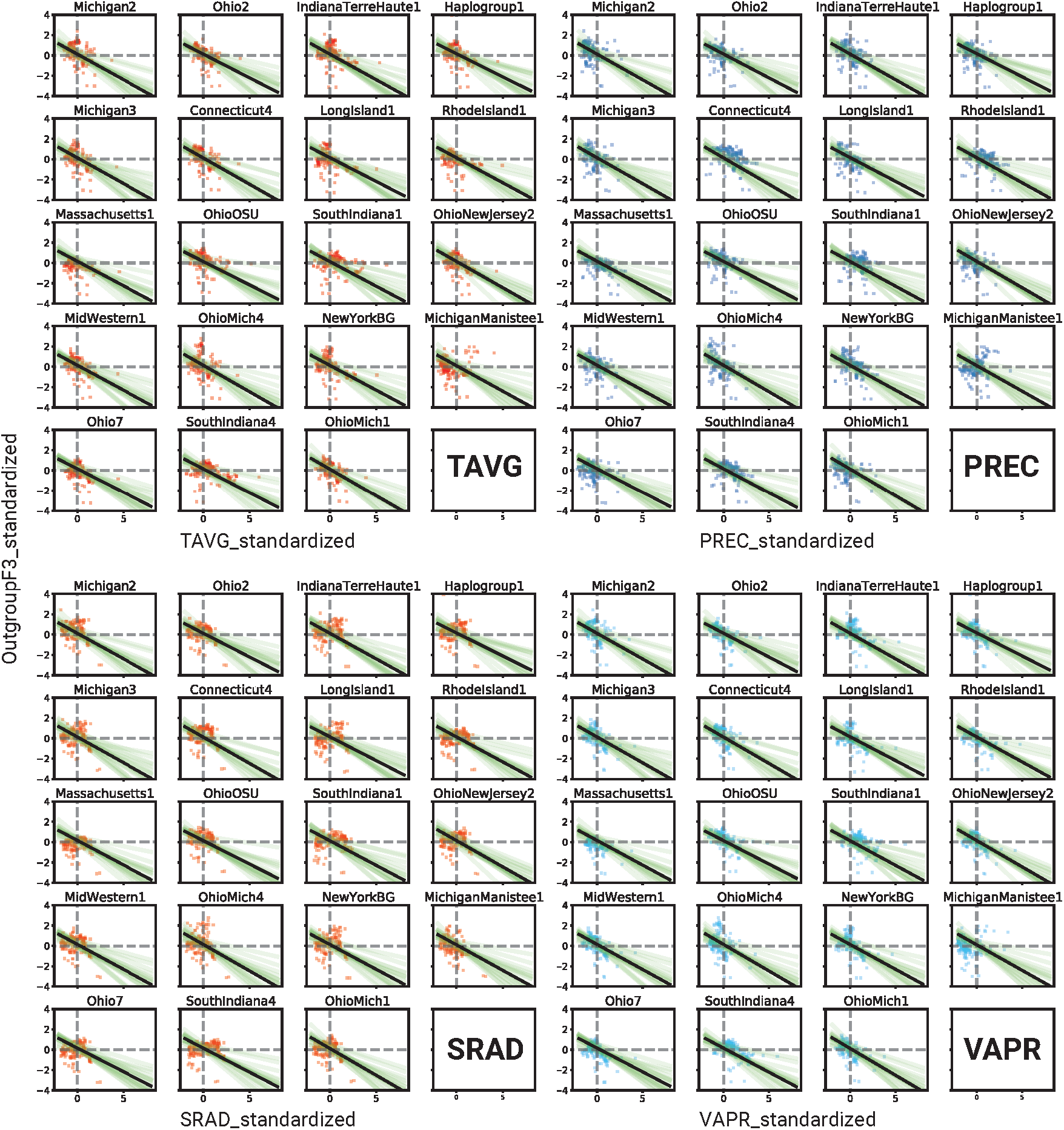
Sampled posterior regression lines for the model describing relationship between outgroup *f*_*3*_-statistics and environmental variables. For every N. American group, mean estimate (solid black line) of the posterior regression lines (thin green lines) was plotted against each environmental variable. X-axis is the standardized environmental variable and y-axis the standardized *f*_*3*_ outgroup statistic derived from the configuration: N. American group, AEA-subcluster_i_: relictsFs12 (as outgroup). The thin green regression lines show overall uncertainty for each group.

**Figure S11.**
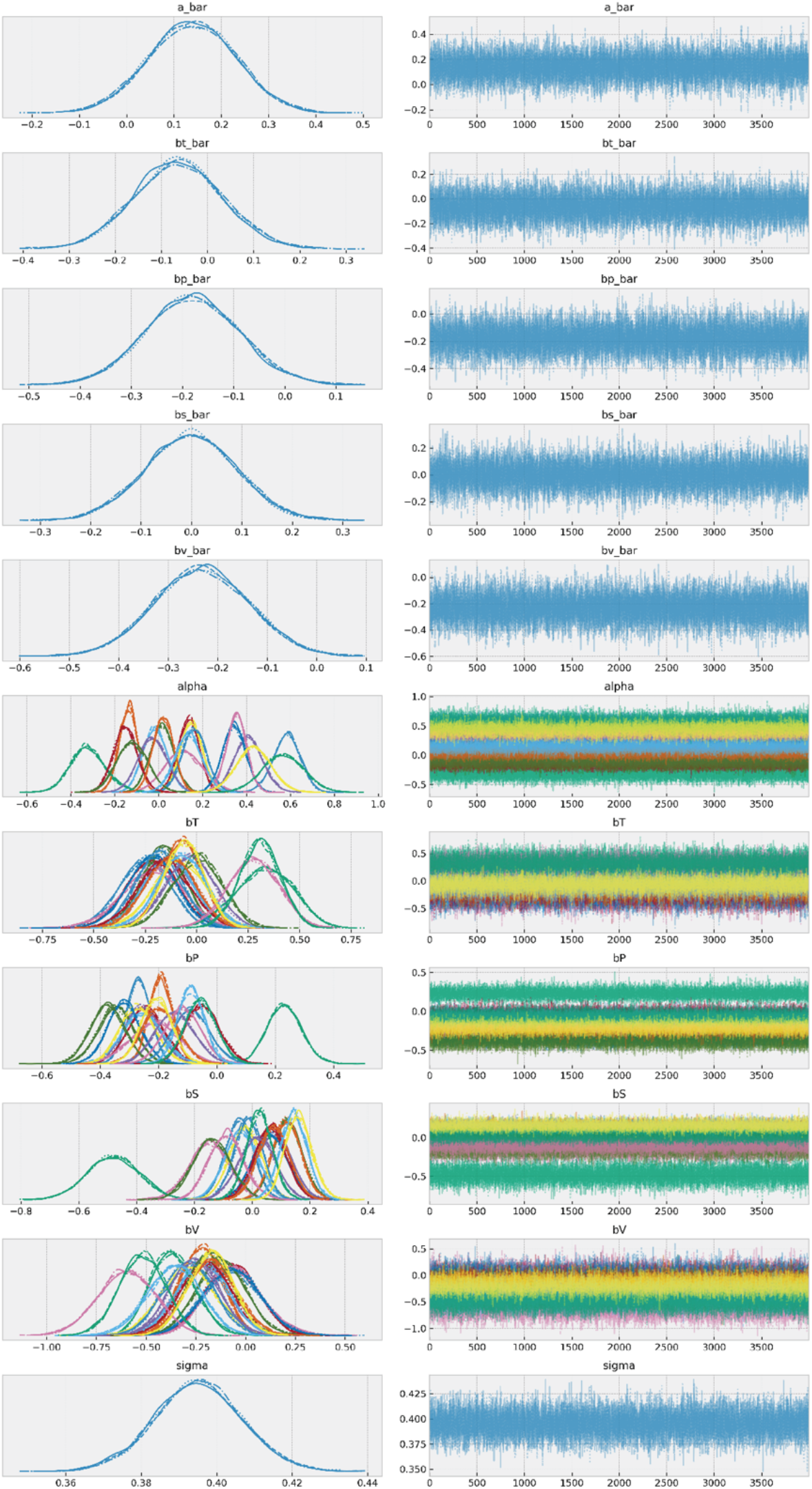
Trace plots for Bayesian Multi-level Model (bMLM) Left columns are marginal values of the trace for different parameters (a_bar = pooled intercept, bt_bar = pooled *β* coefficient for tavg (°C), bp_bar = pooled *β* coefficient for precipitation (mm), bs_bar = pooled *β* coefficient for solar radiation (kJ m^−2^ day^−1^), bv_bar = *β* coefficient for water vapor pressure (kPa), alpha, bT, bP, bS and bT are the respective individual group’s *β* coefficients). Right columns are the model traces from 4,000 iterations after 1,000 tuning iterations for the parameters.

**Figure S12.**
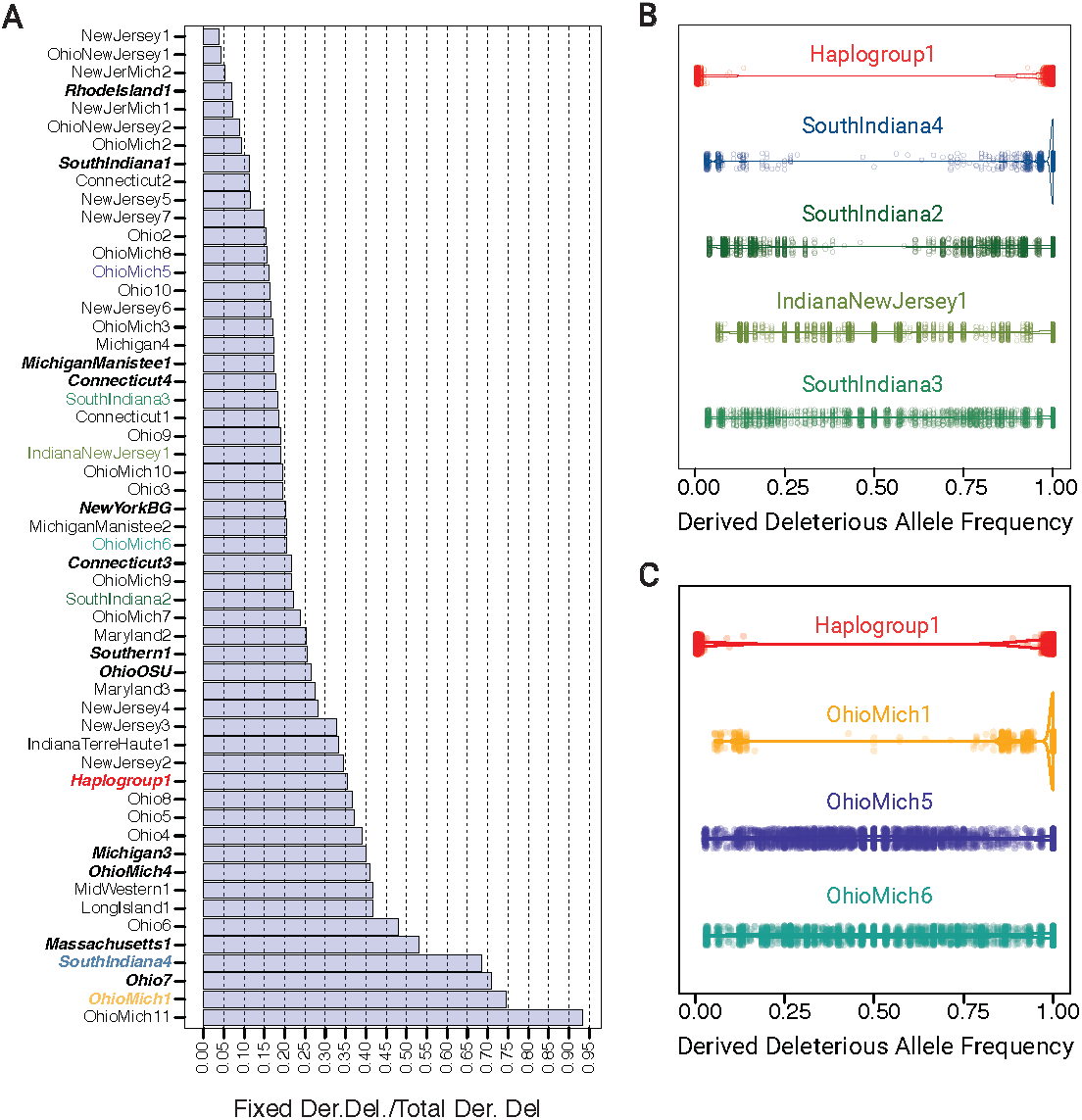
Derived deleterious allele frequencies in admixed haplogroups. **A.** Proportion of fixed derived deleterious to total derived deleterious SNPs in N. American haplogroups. **B.** Derived deleterious allele frequency in haplogroups from Indiana (admixture between Haplogroup1 and SouthIndiana4). **C.** Derived deleterious allele frequency in haplogroups from Ohio/Michigan (Admixture between Hap-logroup1 and OhioMich1).

